# Nuclear Calprotectin mediates Epithelial Wound Healing Defects in Crohn’s Disease related Fistula

**DOI:** 10.1101/2025.03.06.641817

**Authors:** Marte A.J. Becker, Pim J. Koelink, Sarah Ouahoud, Sander M. Meisner, Manon van Roest, Vanesa Muncan, Andreas P. Frei, Gerald Nabozny, Frank Li, Job Saris, Dalia A. Lartey, Willem A. Bemelman, Geert R. D’Haens, Christianne J. Buskens, Manon E. Wildenberg

## Abstract

Wound healing is critical to homeostasis in particular in tissues exposed to environmental insults such as the skin and the intestine. At the intersection of these tissues a particular form of wounding occurs in the form of perianal fistula, i.e. abnormal tracts connecting the intestine with the skin. Importantly, a natural disparity is found in healing efficiency in perianal fistula depending on the underlying medical condition, which is currently not well understood. Using this disparity, we modelled appropriate and defective wound healing by comparing Crohn’s disease related fistula (poor healing) and non-IBD/idiopathic fistula (well healing). We show that during wound healing, cytokines TNFα and IL6 in conjunction with TGF-β induce differentiation of columnar intestinal mucosa into squamous epithelium with high expression of keratins 5 and 13 and *WFDC2*. Specifically in poorly healing wounds, cytokines including IL17, IL22 and IFNγ induced expression of S100A8/9 in this intestinal derived tissue. Surprisingly, S100A8/9 was not released but rather functioned as a transcriptional (co)regulator, affecting downstream inflammatory mediators such as C3 and CXCL17. These data identify separate immune pathways contributing to healing in general and deficient healing, allowing for more targeted intervention approaches.

## INTRODUCTION

Wound healing and tissue regeneration are important components in maintaining functional homeostasis in virtually any organism. In particular tissues which are frequently exposed to environmental insults such as the skin and the intestinal mucosa require exceptional healing and regenerative capacity. Failure to heal may result in a variety of pathological conditions, including chronic inflammatory diseases (inflammatory bowel disease, psoriasis), increased infectious risk and chronic wounds. The perianal region is an intersection of skin and intestinal mucosa and is prone to a specific form of pathology, perianal fistula. Fistula are abnormal tracts connecting two epithelial surfaces, in this case the rectum and the skin. Although perianal fistula may occur spontaneously in otherwise healthy individuals (then termed idiopathic or cryptoglandular fistula), they are particularly common in patients suffering from Crohn’s disease. Crohn’s disease is an inflammatory bowel disorder characterized primarily by ulcerated lesions throughout the intestinal tract. However, a substantial proportion (up to 30%) of patients develops additional complications in the form of fistula, causing fecal incontinence, recurrent abscesses and pain, and profoundly impacting quality of life.(1)

Formation of a perianal fistula is a physiological reaction to mucosal wounding in the rectum, in particular in response to local abscess formation. Due to the high local pressure in the abscess, a tract is forced towards nearby epithelial surfaces, allowing drainage and initiating a wound healing response. In patients suffering from idiopathic fistula, this responses is fairly effective, resulting in creation of a stable fibrotic or epithelized tract, amenable to surgical closure with response rates around 81%.(2) In contrast, Crohn’s disease related fistula are highly refractory to therapy, with healing rates usually not exceeding 30-50%, even using a combination of medical and surgical treatment and patients are at high risk for proctectomy or fecal diversion (ostomy).(3) (4)

The biology underlying the difference in healing efficacy between Crohn-related and idiopathic fistula remains incompletely understood. Interestingly, neither the occurrence nor the refractory nature of Crohn-related fistula are correlated directly to the luminal activity of the disease, suggesting differential pathogenesis for fistula and luminal inflammation. Several studies have suggested epithelial to mesenchymal transition (EMT) as a contributor to fistula formation and as culprit in the pathology (5, 6), primarily based on the expression of EMT related proteins including TGF-β, SNAIL and ZEB within fistula tracts. However, expression of EMT related factors was also described in idiopathic fistula, indicating it may be more related to the general wound healing response than have a role specifically in the disturbed healing in Crohn’s disease.(7)

One factor complicating studies into the biology of Crohn-related fistula is the known heterogeneity within the patient group. Although on average Crohn-related fistula display less efficient healing, this varies considerable between patients, creating mixed samples. Recently, a new fistula classification was implemented, which stratifies Crohn’s disease related fistula strongly taking into account the likelihood of fistula closure (i.e. wound healing).(8) This stratification allows for generation of more clinically homogeneous patient groups, thus improving resolution of the data. In this study, we assessed perianal wound healing responses by systematically comparing severe and mild Crohn-related fistula (poor and intermediate healing respectively) to fistula (good healing).

We identify a wound healing program which involves re-differentiation of mucosal epithelium into squamous stratified epithelium expressing specific keratins, including KRT5 and KRT13. The redifferentiation involves an intermediate tissue type expressing WFDC2, likely induced by the inflammatory process. In Crohn-related fistula, the wound healing response was derailed by nuclear effects of S100A8/9 (calprotectin). When expressed in the intermediate WFDC2+ epithelium, nuclear expression of S100A8/9 functioned as a transcriptional (co)regulator promoting inflammation and resulted in generation of disorganized and inflammatory tissue.

These data reveal a new wound healing program occurring at the mucosal-skin interface and identifies epithelial calprotectin as a dysregulating factor in wound healing.

## RESULTS

### Identification of a keratinization program as a general mucosal wound healing response

Given the difference in therapy responsiveness and Crohn-related fistula transcriptional profiles of fistula tracts were compared between both groups. To avoid variation due to fistula location, only samples obtained specifically from perianal fistula were included. Overall, 136 differentially expressed genes were identified (FDR<0.05, Fig 1a). Pathway analysis revealed an increased activation of immune responses, in particular adaptive immune responses, in the Crohn-related fistula, in line with the underlying immunological condition. In contrast, fistula showed increased activity of pathways associated with (epi)dermal differentiation, such as *keratinization* and *epidermis development* (Fig 1b). Potential confounders in this analysis include Crohn’s specific medication (e.g. anti-TNF), the presence of a deviating colostomy (predominantly in Crohn’s patients) and a longer fistula duration in Crohn’s patients. However, correction for these factors did not affect the results, neither did the presence of a seton (Fig S1).

**Figure 1.**
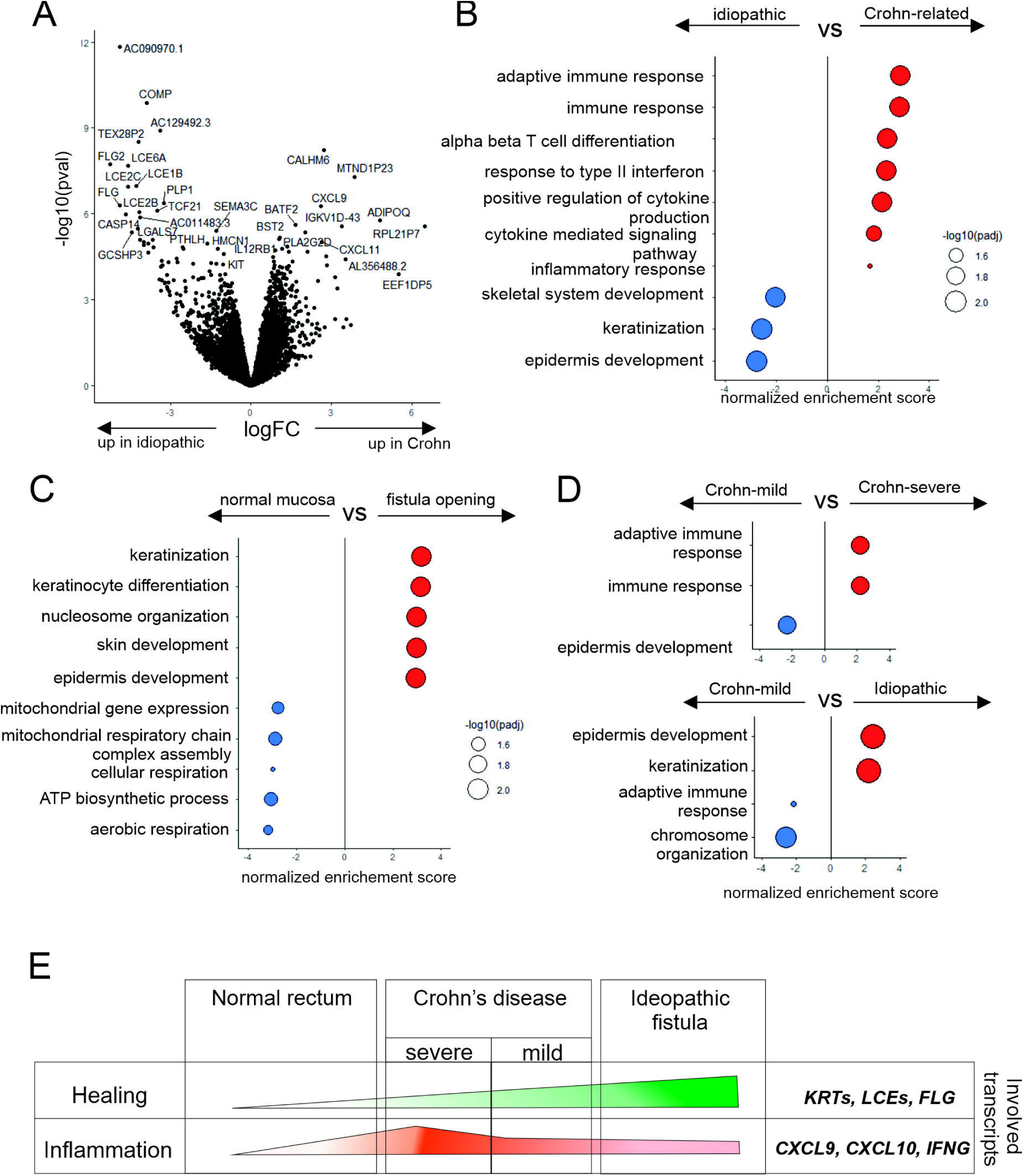
BulkRNA Seq analysis of fistula tracts of peri-anal fistula indicates epidermal differentiation as a wound healing response. (A) Samples of Crohn’s related (n=38) and non-IBD/idiopathic fistula (n=13) were obtained and gene expression profiles generated using bulk RNASeq. (B) Gene Ontology pathway enrichment analysis of the expression profiles of Crohn’s related vs idiopathic perianal fistula. (C) Biopsies were obtained from the internal opening of Crohn-related peri-anal fistula (n=20) and compared to RNASeq data for non-inflamed rectum obtained from two public resources (GSE117993, n=48 and GSE83245, n=21). (D) Gene Ontology pathway enrichment analysis of the transcriptome of severe vs mild Crohn fistula (top) and mild Crohn-related versus idiopathic fistula (bottom). (E) Schematic overview of differences between normal rectum, mild and serve Crohn-related fistula and non-IBD/idiopathic fistula.

As comparative analyses provide relative activity levels, the next question was whether the keratinization response observed was completely absent in Crohn-related fistula or just decreased compared to idiopathic fistula. To this end, transcriptional profiles of the internal fistula opening were compared to those obtained from normal rectal mucosa. As no wounding has occurred in normal mucosa, wound healing activity is absent here. Strikingly, the most differentially active pathways were again those associated with (epi)dermal differentiation which were significantly more activated in the Crohn-related fistula (Fig 1c). This supported a decrease rather than complete absence of keratinization in Crohn related fistula. Clinically it has been well recognized that not all Crohn’s fistula behave similarly, and a classification system was recently established based on disease severity and likelihood of fistula closure. Applying this classification we compared mild (closure more likely, class 2a/b) to severe (requirement of ostomy or colectomy, class 2c/3) Crohn-related fistula. Pathway analyses indicated that the keratinization/epidermal differentiation response was most pronounced in fistula, decreased in mild Crohn related fistula, even more decreased in severe Crohn fistula and absent in normal mucosa (Fig 1d/e). In summary, activity of the keratinization/epidermal differentiation was correlated with the efficacy of fistula healing.

### The wound healing associated ‘epidermal differentiation’ includes an intermediate tissue with characteristics of both mucosa and squamous epithelium

We then aimed to further delineate the keratinization wound healing response we identified. Histological evaluation of fistula biopsies showed the presence of stratified squamous epithelium, in line with the ‘epidermal development’ and ‘keratinization’ identified (Fig 2a). In addition, columnar rectal mucosal epithelium was observed in most samples. Interestingly, in several samples the tissue adjacent to the squamous regions was less structured and cuboid, suggesting an intermediate tissue phenotype between columnar and squamous epithelium.

**Figure 2.**
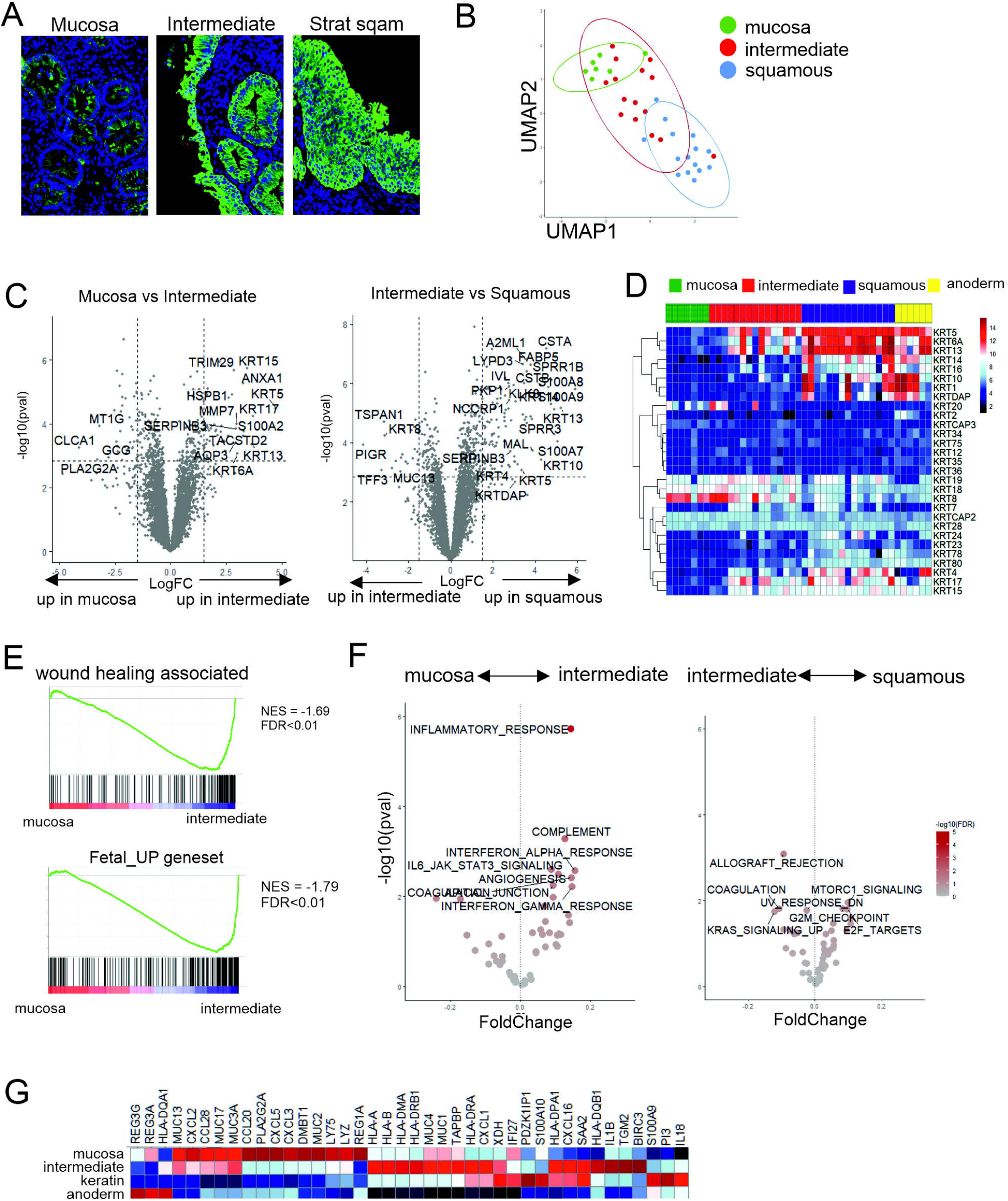
Digital spatial profiling of mucosal, intermediate and squamous epithelium in fistula openings. (A) Representative image of the three types of epithelium selected, stained for pancyokeratin (green) and DAPI (blue). (B) DSP analysis was performed on the mucosal (n=7), intermediate (n=15) and squamous tissue (n=15) in the internal fistula opening using a whole genome library. Clustering by UMAP is shown. (C). Differential expression analysis of the transcriptional profiles obtained with FDR multiple testing correction. (D) Heatmap of expression of keratin transcripts. (E) Geneset Enrichment analysis of genesets previously described to be associated with wound healing associated epithelium or increased in fetal intestine compared to adult. Normalized enrichment score is shown. (F) Pathway analysis on transcriptional profiles of indicated tissues using the Hallmark pathway dataset. (G) Heatmap depicting per sample expression of transcripts previously associated with ‘LDN’ epithelium in the intestine.

To better define these epithelial tissue subtypes, we performed digital spatial profiling (DSP), analyzing normal appearing mucosa, stratified squamous epithelium and the intermediate appearing regions in fistula openings. Dimensionality reduction of samples showed clear separate clustering of the regions defined as normal appearing mucosa” and “stratified squamous epithelium” (Fig 2b). The cuboid ‘intermediate’ tissue clustered in between the other two subsets, supporting not only a morphologic but also transcriptionally intermediate phenotype. For example, the keratin expression profile of the intermediate cells was a combination of the keratins expressed by the intestinal mucosa (e.g. *KRT8, KRT18*) and those expressed in the squamous tissue (e.g. *KRT5, KRT13*). Typical epidermal associated markers such as *KRT1* and *KRT10* were expressed in the squamous areas, however not to the same extend as in the anoderm, which was included for reference purposes (Fig 2c/d). The intermediate phenotype was characterized by expression of *WFDC2*. Although some markers associated with EMT indeed were expressed in the internal openings (*TGFB1, SNAI2, ITGB6, TWIST1,* Fig S2a) as previously described by others, the total EMT program was not specifically activated in any of the tissue subtypes (Fig S2b).

Re-differentiation of the intestinal mucosa is a form of plasticity which has previously been described in murine models, and was then referred to as ‘wound associated epithelium’ or ‘fetal-like reprogramming’.(9, 10) Geneset enrichment analysis indicated that the intermediate tissue indeed shows significant enrichment for both fetal and wound healing associated programs (Fig 2e). The previous studies suggested a role for the inflammatory response in initiation of this process, which was also supported by pathway analysis in our data, as most pathways differing between normal mucosa and intermediate tissue were related to inflammation (e.g. Hallmark pathways ‘*Inflammatory response’*, ‘*Interferon signaling’*, and *‘IL6-Jak-Stat3-signaling’*, Fig 2f).

Recently, a human study described a subset of epithelial cells nearly exclusively present in IBD patients, which is characterized by expression of LCN2, NOS2 and DUOX2. Although most of the genes associated with the ‘LND signature’ were also present in our study, this was not specifically enriched in any specific tissue type (fig 2g).

### Human adult mucosal tissue can develop into the intermediate WFDC2+ tissue phenotype

Both the morphology and the transcriptional profile of the intermediate tissue suggested a transitional state between the mucosa and the squamous epithelium. To experimentally validate the potential of intestinal epithelium to differentiate into the intermediate and keratinized tissue type, an in vitro model was employed. Upstream regulator prediction using the DSP data suggested that the responses observed in the intermediate tissue was consistent with activity of TNF-α, TGF-β and IL6 (Fig 3a). Organoid cultures were established from adult colon mucosa and stimulated using either a mixture of TNF-α, TGF-β and IL6, or TGF-β alone. Stimulation with the cytokine mix resulted in a significantly higher number of organoids with a squamous appearance, in line with the phenotype observed in vivo (Fig 3b). TGF-β alone did not have similar effects. Analysis of specific markers that characterized the three tissue subtypes in our DSP analyses (Fig 3c, left) showed a strong increase of intermediate transcripts *MMP7* and *WFDC2* and a moderate increase in squamous tissue associated transcripts *CALML5* and *KRT13* although the last failed to reach significance (Fig 3c, right). Interestingly, the decrease in mucosal associated transcripts was most pronounced by stimulation by TGF-β alone, while the increased expression of intermediate and squamous markers was significantly stronger upon stimulation using the cytokine mixture. This may indicate the loss of mucosal markers mainly depends on TGF-β, while the induction of intermediate and squamous transcripts requires additional cytokine signaling.

**Figure 3.**
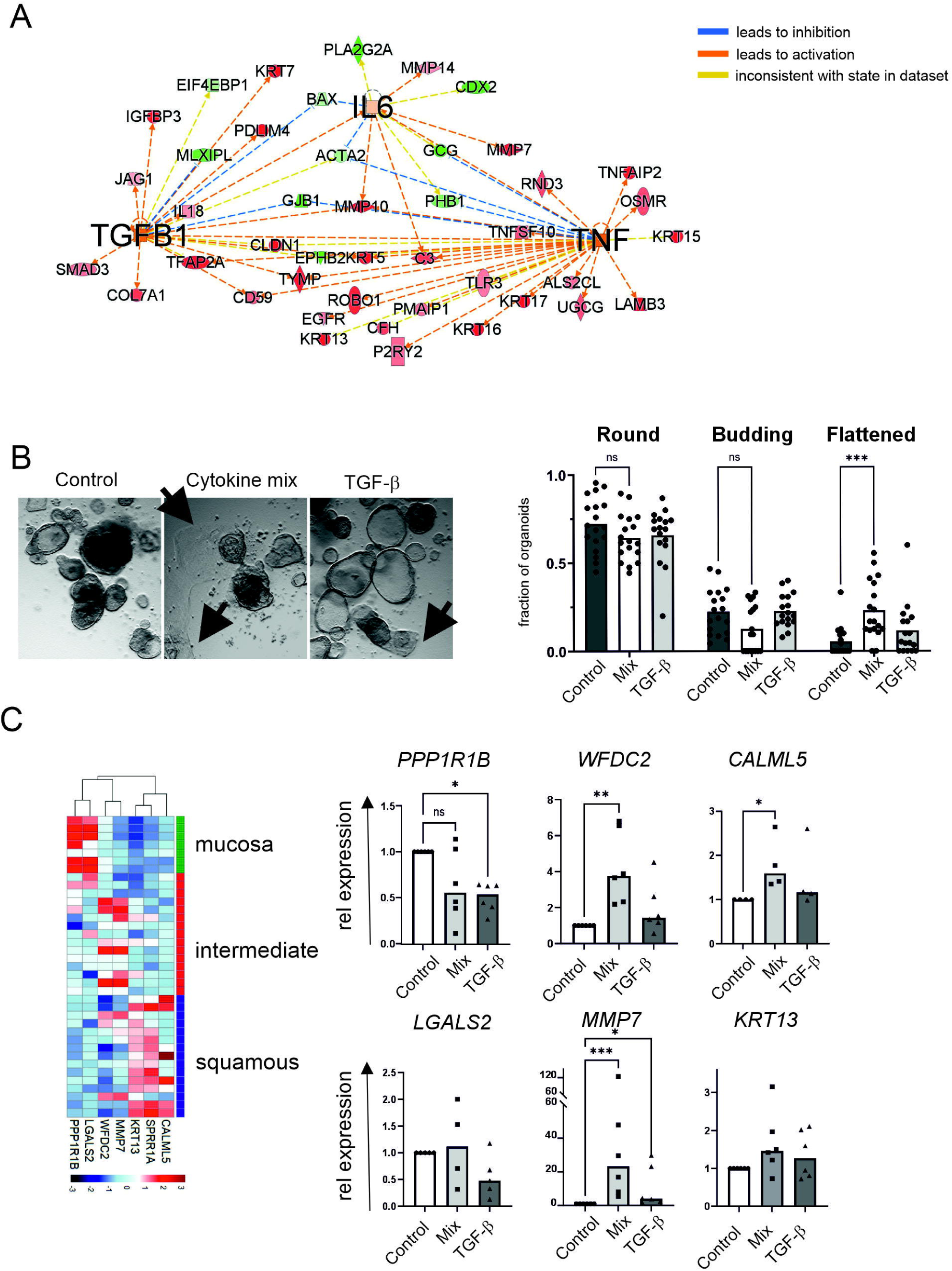
Adult human organoid culture displays transcriptional profile similar to that seen in wound healing. (A) Digital Spatial Profiling data was used to predict upstream activity in the intermediate tissue type using ingenuity pathway analysis. (B) Human organoids were obtained from adult donors, and stimulated using a cytokine mixture (TNF-a,10ng/mL, TGF-b, 2ng/mL and IL-6, 2ng/mL) or TGF-b alone (2ng/mL). Subsequently, images were obtained and analyzed for the presence of rounded, budding or flattened organoids (n>250 cells per condition, 2 individual donors). Bars represent median, dots represent individual images., two-way Anova, corrected for multiple testing by Dunnets. *** denotes p<0.0001. (C) Left panel displays DSP expression profile of the genes selected for organoid culture analysis. Right panel depicts quantitative pcr results shown as relative to the control of the same donor (n=4-6). Bars represent median, dots represent individual experiments of independent donors. Statistics was performed using Kruskall Wallis test with Dunn’s multiple correction, * denotes p<0.05, ** p<0.01, *** p<0.001.

### Aberrant development of squamous epithelium in Crohn’s disease fistula

Clinically, wound healing is clearly impaired in Crohn-related fistula, and the RNA data suggest this is correlated to a decrease in the mucosa-to-keratinized transition. We explored this further using a single cell approach. The epithelial fraction of biopsies obtained from the fistula of 20 individual Crohn’s disease patients as well as 10 idiopathic controls were analyzed. Clustering of epithelial cells showed seven major clusters, including enterocytes, Goblet cells, Tuft cells and enteroendocrine cells (Fig 4a and Fig S3). One cluster of particular interest in light of our data was characterized by relatively high levels of *WFDC2*, similar to the phenotypically ‘intermediate’ tissue we observed earlier. Keratin expression in the WFDC2+ population in the single cell analysis was also comparable to a mixture of intermediate and squamous tissue (*KRT19, KRT5, KRT13*, Fig 4b).

**Figure 4.**
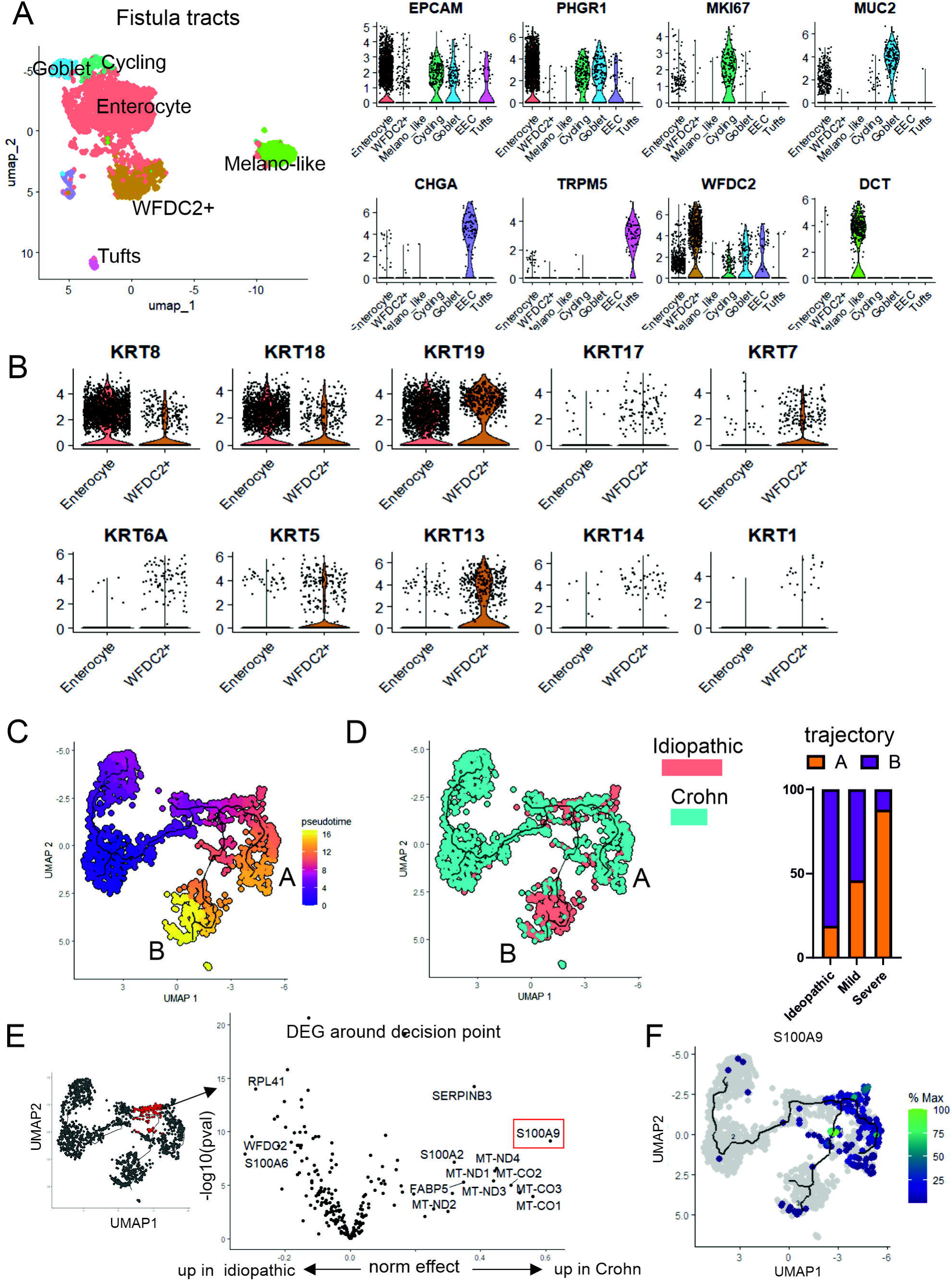
Single cell analysis of the epithelium in internal openings reveals a subset of WFDC2+ cells which is dysregulated in Crohn’s disease. (A) UMAP clustering of the epithelial cells obtained from the internal opening of 31 fistula (idiopathic n=10, Crohn n=20). Violin plots shows overall data as well as expression in individual cells (dots) for cell type characteristic markers.(B) Expression of selected keratins. (C) Pseudotime analysis of the intermediate to late stages of development (based on Fig S4). (D) Proportion of cells locating to either trajectory A or B, separated per fistula subtype. (E) left panel: selection of cells for the evaluation of expression around the branching point between trajectory A and trajectory B. right panel: differential expression between idiopathic and Crohn for the cells selected in left panel. (F) Expression of S100A9 over the developing WFDC2+ epithelium.

Using the single cell dataset, we were able to model the development of WFDC2+ subsets by pseudotime analysis. The results suggested a development from *PIGR* and *PLA2G2A* expressing enterocytes into the WFDC2+ expressing subsets (Fig S4b). Notably, differentiation appeared to occur in two separate trajectories (denoted trajectory A and B, Fig 4c). Pathway analysis of these two trajectories showed pro-inflammatory activity in trajectory A with enriched pathways such as ‘*Response to bacterium’, ‘Antigen processing’* and *‘Immune response’* (Fig S4c). Strikingly, distribution of cells over the two trajectories varied strongly between fistula types and correlated with healing efficiency (Fig 4d). The majority of cells derived from fistula were found in trajectory B, while cells from severe Crohn-related fistula were in trajectory A. Cells derived from mild Crohn-related fistula were distributed equally over the two trajectories. In order to identify factors contributing to the altered differentiation path in Crohn’s related fistula, cells around the branching point were compared specifically between Crohn’s and samples. Surprisingly, *S100A9*, one of the subunits of calprotectin was one of the most apparent genes increased in Crohn’s derived early WFDC2+ cells (Fig 4e/f), indicating a potential role in the altered development of this population.

### S100A9 is induced in CD fistula associated epithelium and acts as a pro-inflammatory transcriptional co-activator

It is generally assumed that in the (inflamed) intestine, calprotectin is derived from infiltrating neutrophils and not expressed by the epithelium. To assess whether the signal obtained in this study was truly epithelial expression of S100A9, sections of the internal openings of both Crohn and idiopathic fistula were stained by immunohistochemistry. Indeed, expression of S100A9 was observed in the more squamous and intermediate appearing mucosa, but not in adjacent normal intestinal mucosa (Fig 5a and Fig S5). Strikingly, expression was partially nuclear, in particular in Crohn’s derived samples.

**Figure 5.**
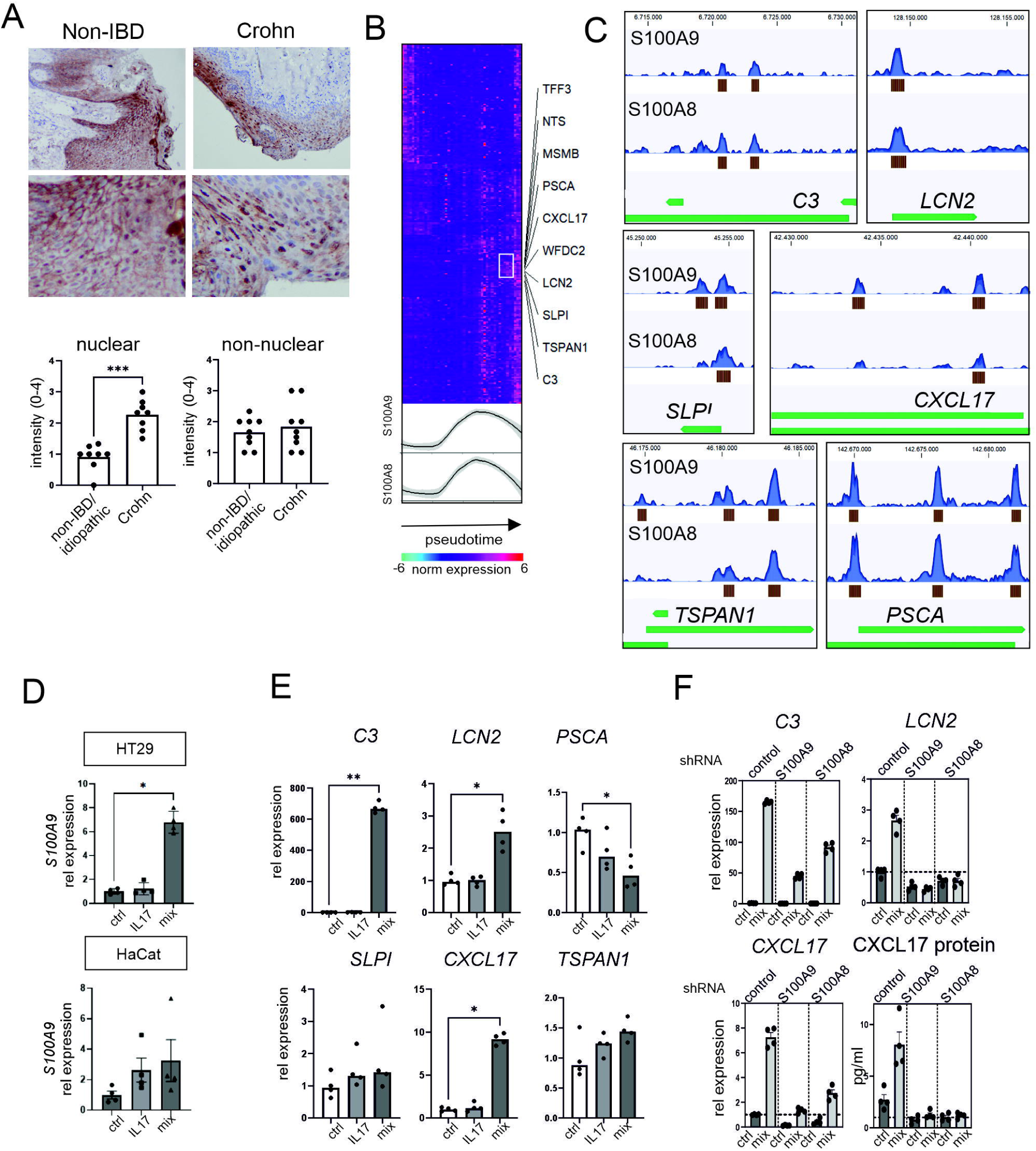
Nuclear expression of calprotectin is increased in squamous epithelium in Crohn’s disease and may serve as a pro-inflammatory transcriptional coregulator. (A) Immunohistochemical staining of S100A9 in the internal openings of idiopathic (n=9) and Crohn (n=9) related fistula. Bottom panel: Quantification of nuclear and non-nuclear staining on a scale 0-4. Bars represent mean, dots represent individual slides. Kruskall-Wallis with Dunn’s multiple testing correction, *** p<0.001. (B) Heatmap representing the expression over pseudotime of all genes differentiating trajectory A and B in figure 4E, lines representing expression of S100A8 and S100A9 over pseudotime. (C) Reanalysis of ChipSeq data using S100A9-Flag or S100A8-Flag as IP target in transformed MCF-10A–ER-Src cells. Top row shows sequences identified in S100A9 precipitation, middle row S100A8 precipitation, bottom rows show localization of the relevant gene, brown blocks denote significant peaks (p<0.05). (D/E) Intestinal cells (HT29) were stimulated using a mixture of cytokines (IL17 50 ng/ml, IL22 50 ng/ml, IFN-γ 10 ng/ml, TNF-α 20 ng/ml) or IL17 alone for 24 hours. RNA expression of is shown as relative to the unstimulated cells. Bars represent mean, Mann-Whitney U-test, * p<0.05. (F) S100A8 or S100A9 where knocked down using lentiviral shRNA and cells were stimulated as in D. Bottom right: cells were stimulated for 72h and supernatants analyzed for CXCL17 protein by ELISA.

The nuclear localization of S100A9 suggests a transcriptional role for calprotectin in these cells, which has also been suggested in earlier studies.(11, 12) The pseudotime analysis allowed us to compare the expression profiles of genes differentially expressed between the two trajectories observed above, relative to the timing of S100A8/9 expression (at the branching point). A specific cluster of genes (*TFF3, TSPAN1, MSMB, NTS, PSCA, CXCL17, WFDC2, LCN2, SLPI, C3*) was upregulated closely after the peak increase in *S100A8/9* expression in pseudotime (Fig 5b). We then reanalyzed a publicly available CHIP dataset with a focus on the genes upregulated close after *S100A8/9* in pseudotime. Indeed, peaks were observed near the transcription start sites for 6/10 of these genes *(C3, LCN2, SLPI, CXCL17, TSPAN1, PSCA*), indicating potential transcriptional regulation by S100A8/9 (Fig 5c).

Data regarding epithelial expression and function of S100A8/9 in the intestine is scarce, but expression and regulation has been well documented in psoriatic skin lesions where expression of S100A8/9 was maintained through interaction with the IL17/IL23 axis.(12, 13) Interestingly, in a keratinocyte cell line, IL17 alone was shown to be sufficient to induce expression of S100A9, while the intestinal epithelial cell lines required a mixture of cytokines including IL17.(13, 14) We confirmed this data, showing an inflammatory cytokine mix containing IL17 (TNF-α, IFN-γ, IL22 and IL17), but not IL17 alone induced *S100A9* in HT29 intestinal cells (Fig 5d). In contrast, and in line with the data from the psoriasis field, IL17 alone was sufficient for induction in skin derived HaCat cells (Fig S6a). In three additional intestinal epithelial cell lines (Caco2, HCT116 and T84) expression of *S100A9* was also increased upon stimulation with the cytokine mixture (Fig S6b). Out of the six previously identified potential transcriptional targets, *C3, CXCL17* and *LCN2* indeed increased upon stimulation and S100A9 induction (Fig 5e). Binding of S100A8 and S100A9 to the relevant sites in the C3, CXCL17 and LCN2 was confirmed by CHIP-pcr on intestinal epithelial cells (Fig S6c). Supporting a critical role of S100A9 in the induction of the downstream targets, knockdown of either S100A9 or S100A8 resulted in decreased or abrogated upregulation of *C3, CXCL17* and *LCN2* transcripts as well as abrogated induction of CXCL17 protein secretion (Fig 5f). As extracellular S100A8/9 dimer can also function as a pro-inflammatory cytokine, we assessed secretion of calprotectin. Similar to previously published data, extracellular S100A8/9 could not be detected after stimulation of intestinal epithelial cells (data not shown). (14) In contrast, immunoprecipitation of nuclear S100A8/9 did show co-precipitation of NFkB and STAT3, transcription factors downstream of IL22, IL17 and TNF-α signaling, suggesting a role for nuclear calprotectin as a transcriptional co-factor (Fig S6d). (14) Together, this data suggests calprotectin as a critical regulator of various downstream targets upon stimulation of wound healing tissue with a pro-inflammatory cytokine mix mimicking the fistula environment.

### Primary human intestinal epithelium and tissue express nuclear S100A8/9 and downstream CXCL17

Finally, to validate these findings in primary intestinal cells, adult colon organoids were stimulated using the inflammatory cytokine cocktail (TNF-α, IFN-γ, IL22 and IL17), and also showed strong upregulated expression of both S100A8/9 as well as putative targets *C3, CXCL17* and *LCN2*. Importantly, expression of S100A8/9 peaked early in stimulation, while the putative targets continued to increase (Fig 6a). To assess the physiological relevance in Crohn’s disease, we analyzed expression of *CXCL17* mRNA in tissue sections of the internal fistula openings. *CXCL17* was clearly expressed in the squamous epithelial tissue, in particular in areas also expressing S100A9 (Fig 6b). Quantification showed increased *CXCL17* expression in Crohn’s disease fistula compared to idiopathic fistula, consistent with the earlier described increase in (nuclear) S100A9 (Fig 6c).

**Figure 6.**
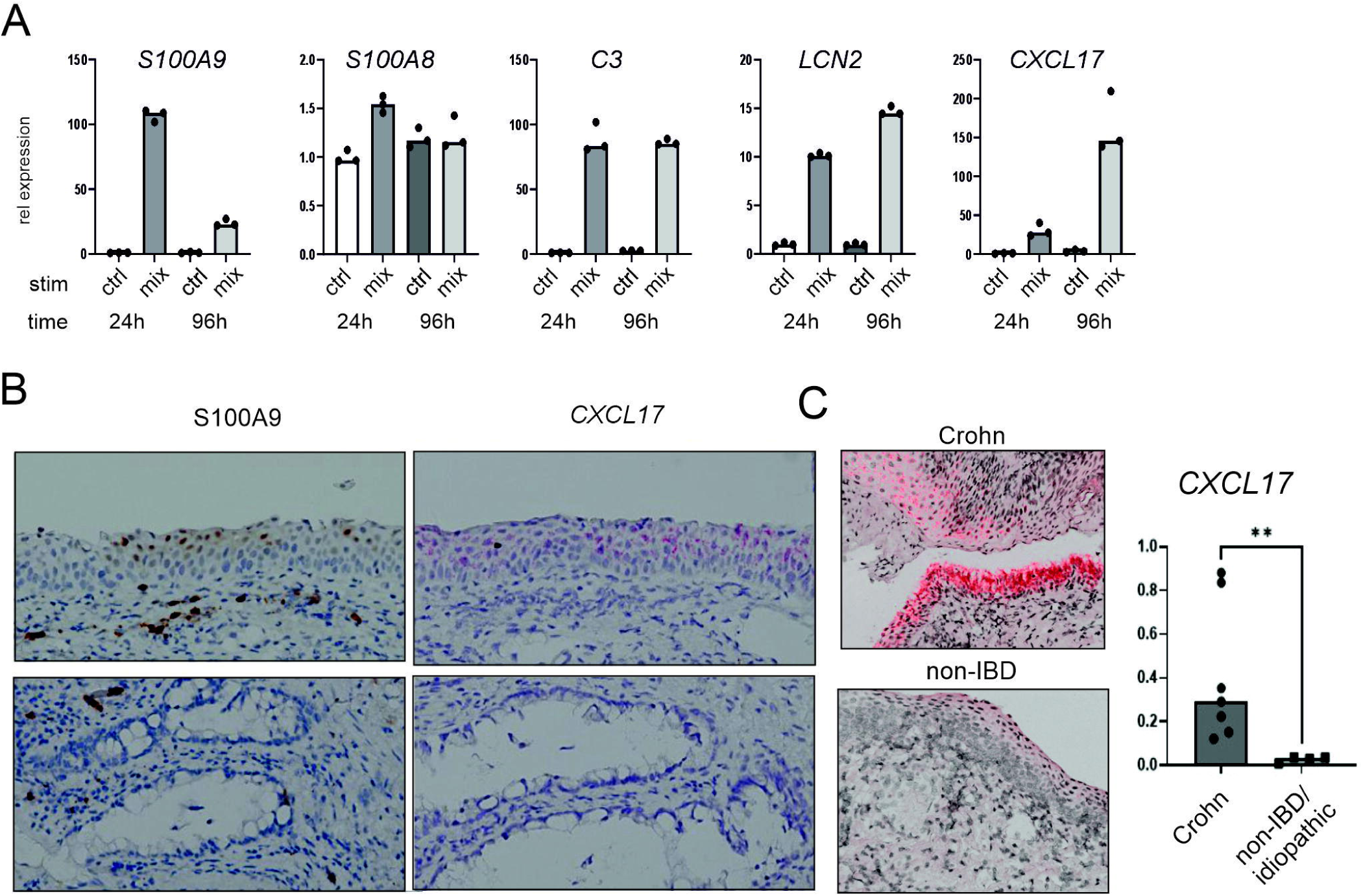
Expression of S100A8/9 and putative targets in patient derived samples. (A) Expression of S100A8, S100A9 and putative downstream target mRNA in human colon organoids treated with cytokine mix for 24h and 96h. Expression normalized to housekeeping genes. (B) Immunohistochemical staining for S100A9 (left, brown) and in situ for *CXCL17* (right, red) on sequential sections indicating co-localization in squamous epithelium. (C) In situ for *CXCL17* (red) on the internal openings of Crohn (n=7) and idiopathic (n=4) related fistula. Representative staining and full quantification shown. Bars represent median, Mann-Whitney U-test, ** p<0.01.

## DISCUSSION

In this study, we show that mucosal wound healing of the rectum involves the development of squamous epithelium expressing specific keratins (*KRT5, KRT13*). In fistula associated with Crohn’s disease, this process is initiated to a certain degree, correlating to the classification of the fistula and thus healing perspective. However, in particular in Crohn’s disease related fistula with a severe phenytp, but diverges to an inflammatory state features mediated by epithelial S100A8/9 (calprotectin).

We show here that upon fistula formation a wound healing program is initiated which results in the development of stratified squamous epithelium rather than the typical columnar intestinal epithelium. The origin of this tissue is an intriguing topic, with two major, non-mutually exclusive, possibilities. Earlier studies showed development of squamous epithelium in mouse models of rectal damage. (15, 16) In these studies, authors describe a subset of anorectal transitional zone cells present in steady state and expanding upon injury, contributing to both squamous and columnar epithelium. Whether the intermediate cells we observe in our study also derive from this ATZ cells remains to be determined. Some markers previously described in ATZ cells were also expressed in the intermediate tissue in our study (*KRT7, KRT17*) although others were not (*APOE, CD34,* data not shown). It should be noted that the previous studies were performed in mouse models, and specific markers for cell types may differ between species. A recent study described the occurrence of LCN2+ epithelial cells in the ascending colon in active Crohn’s disease.(17, 18) This would suggest this type of epithelium also occurs in wounding circumstances in the proximal colon arguing against a pure ATZ derived origin. However, again, expression profiles of these ‘LDN’ cells do not fully overlap with the cells described in this study (Fig S3c). A second possibility is the dedifferentiation or reprogramming of the rectal epithelium. This has also been described in the context of intestinal damage and was termed ‘wound healing associated epithelium’ (WAE) or ‘primitive reprogramming’.(9, 10) When overlaying our data with the gene sets associated with these phenomena, highly significant gene set enrichment was observed, suggesting this may indeed be a contributing factor in the current study.

In earlier studies, epithelium to mesenchymal transitioning (EMT) has been suggested as a major driver of fistula formation, but we did not detect a clear activation of the EMT program.(5, 6) Although this may appear contradictory, it is also a matter of interpretation. Much of the previous work focused on a limited number of EMT associated markers, and showed these were upregulated in fistula when compared to normal mucosa. Indeed, when focusing on individual markers such as decreased *CDH1* (E-Cadherin), or increased *TGFB1* and *ECM1*, we do corroborate these earlier findings. However, mere expression of just a few genes associated with EMT does not constitute the occurrence of the phenomenon of EMT as a whole, which is also supported by the EMT International Association in their guidelines.(19) When evaluating the full arsenal of EMT associated genes, no clear activation could be seen in any of the tissue types in fistula, suggesting the previous studies may have highlighted more of a general wound response (which includes many of the same molecules as EMT) rather than full EMT.

Irrespective of the origin, the wound response described was detected both in and to some degree in Crohn-related fistula. However, single cell analysis revealed altered differentiation of the squamous wound epithelium in Crohn’s disease, with increased activation of inflammatory pathways. Inspection of the gene patterns around the apparent fate decision point indicated calprotectin (S100A8/9) as a critical factor. Fecal calprotectin is widely used in IBD care as a biomarker for intestinal inflammation, and is mainly considered to derive from infiltrating neutrophils. Our data show that while epithelial cells in normal intestinal mucosa do not express S100A9, the intermediate cuboid and keratinized squamous epithelium do. This is not likely to contribute significantly to fecal calprotectin levels, as it is only expressed in a very limited percentage of cells of the rectum, and published results as well as our data suggest the proteins are not secreted from these cells.(14) In contrast, increased intracellular expression was observed in the epithelium of Crohn’s fistula with a particular localization in the nucleus. This localization suggests a transcriptional role, something which has previously been described during transformation of breast cancer cell lines.(11) Re-analysis of public data showed that a number of the genes enhanced in Crohn’s derived squamous tissue is indeed suggested as transcriptionally regulated by S100A9. Subsequent in vitro studies confirmed the critical role of S100A9 in the induction of *C3, CXCL17 and LCN2* in intestinal epithelium by a mixture of inflammatory cytokines including IL17 and IL22.

The downstream effects of this activity may be mixed. From an immunological perspective, upregulation of CXCL17 and LCN2 are described to result in opposing effects, with CXCL17 recruiting myeloid derived suppressor cells and Treg,(20, 21) while LCN2 would mainly increase the recruitment of neutrophils and activated T cells.(22, 23) From an epithelial perspective LCN2 was shown to inhibit proper keratinocyte differentiation resulting in parakeratosis similar to that seen in Crohn’s disease associated fistula.(24) Effects of CXCL17 on keratosis have not been described.

A causative role for S100A8/9 has been described extensively in the context of psoriatic skin lesions.(12, 13) In these studies the IL17/IL23 axis was identified as the major driving force, and IL17 blocking therapy indeed is highly successful in the treatment of patients suffering from psoriasis. (25) (13) Interestingly, a recent study also showed a critical role for IL17 in the development of parakeratosis during aging, and blockade of IL17 was highly effective in reversing skin aging effects in a mouse model. (26) The increased expression of S100A8/9 in the current study was also strongly correlated with the local presence of IL17 producing T cells (data not shown, manuscript in preparation). Previously, IL17 inhibition failed as a therapy for luminal Crohn’s disease due to lack of efficacy of even worsening of disease. (27, 28) However, these studies focused on patients with luminal Crohn’s disease rather than patients with active fistula. In addition, our data suggests that in the intestine, combined IL17 and IL22 effects may be more prominent than the pure IL17 effects seen in the skin. Subsequently, effective therapy in Crohn’s related fistula may require either combined inhibition or interventions in joint downstream mediators. However, due to the multifaceted role of IL22 in the intestine,(29) this will require extensive safety and feasibility evaluation.

In summary, this study demonstrates redifferentiation of mucosa and subsequent development of squamous epithelium as a wound healing response upon the formation of peri-anal fistula. In Crohn’s disease, likely due to the specific immune environment, this process is dysregulated through the transcriptional co-activation activity of calprotectin, resulting in disorganized epithelium less amenable to surgical intervention. Separating general healing from the dysregulation induced by calprotectin may provide leads for future interventions in perianal fistula, supporting healing but inhibiting excessive inflammation.

## Materials and Methods

### Collection of human specimens

Patients suffering from Crohn’s disease related fistula or idiopathic fistula were included in this study. Crohn’s disease was diagnosed based on clinical, biological and endoscopic criteria by the treating physician. Fistula were stratified at the time of sampling according to the TopClass classification.(8) Clinical data was collected regarding gender, age, diagnosis, IBD related parameters and relevant medication (Table S1-S5). Samples were obtained by the treating physician during routine care procedures (inspection under anesthesia or surgery). Included samples consisted of curettage material, biopsy samples of the internal opening of the fistula and the rectal mucosa and full resection specimens. Curettage material was retrieved from within the fistula tract using a curette surgical instrument to create a fresh tract. The internal opening was identified in the course of the procedure (either macroscopically visible or identified by flushing the external opening with saline). The rectal (biopsy) sample was obtained contralateral to the internal opening.

For bulk sequencing, samples were snap frozen and stored at –80°C immediately after collection. For single cell experiments, tissue sections were immersed in fetal calf serum containing 10% DMSO (Sigma-Aldrich), slow frozen using a CoolCell freezing container (Corning) and subsequently stored at –80°C. Full tissue sections were fixed using 4% paraformaldehyde and subsequently embedded into paraffin blocks.

### Ethics

This study involves human participants and was approved by an Ethics Committee or Institutional Board (Biobank review committee of the Academic Medical Centre Amsterdam, (number 178#A201470)). All participants provided written informed consent prior to taking part in the study.

### Organoid cultures

For organoid cultures, colonic biopsies or resection specimens were obtained from 6 adult Crohn’s disease patients (5x non-fistulizing, 1x peri-anal fistula). Samples were washed and the mucosal layer was stripped from the underlying layers. Samples were cut into 5-mm pieces and washed with cold phosphate-buffered saline (PBS) several times until the supernatant was clear. The tissue was incubated in dissociation mix (5 mM EDTA, 2mM DTT, 1% FCS in Advanced DMEM/F12, 4°C, 30 minutes). After incubation, pieces were vigourously mixed using a 25ml pipette, washed in PBS, filtered through a 70 micrometer cell strainer and resuspended in DMEM containing GlutaMAX (Invitrogen), supplemented with penicillin/streptomycin (Gibco) and Hepes (Sigma). Cells were then pelleted, mixed with Matrigel (BD/Corning) and plated in 24-well plates. Isolated colon organoids were cultured in IntestiCult human OGM medium (Stemcell) with 1% penicillin/streptomycin. IntestiCult medium was changed each three days and cultured organoids were passaged and expanded every seven days.

### Organoid differentiation and stimulations

Isolated colon organoids were passaged and allowed to grow out for 3 days. After three days, medium was replaced as described below. For differentiation analyses, medium was replaced by IntestiCult medium containing a cytokine mixture (TNF-α (10ng/mL, Peprotech, 300-01A), TGF-β (2ng/mL, Peprotech, cat 100-21) and IL-6 (2ng/mL Peprotech, cat: 200-06). Medium and cytokines were refreshed every 3 days. For two donors, images were obtained using an inverted Leica DMi8 Microscope (n=10 images per donor per condition). Image analysis was performed by a blinded observer scoring individual organoids as round/cystic, budded or flat. For S100A8/9 induction analyses medium was replaced by Intesticult containing a cytokine mixture (50 ng/ml rhIL17 (R&D,7955-IL/CF Lot DCVA1123021), 50 ng/ml rhIL22 (Peprotech, 200-22, Lot#0710246 I1422), 10 ng/ml rhIFNγ (R&D, 285-IF-100), 20 ng/ml rhTNF-α (Peprotech, #300-01-A Lot #031825 A2323). RNA was extracted using the Bioline Isolate II RNA Mini Kit (Qiagen) according to manufacturer’s protocol.

### Cell lines

HT29, T84, Caco2, HCT116 and Hacat cell lines were obtained from the ATCC and cultured in DMEM (Dulbecco’s modified eagle medium (Lonza, Leusden, The Netherlands), 10% fetal calf serum (FCS), 1 % penicillin/streptomycin (Invitrogen, Thermo Fisher Scientific, Waltham, MA, USA) and 5 mM glutamine (Lonza) at 37 degrees and 5% CO2. Cells were routinely tested for mycoplasma contamination and all tested negative.

Knockdown cell lines for S100A8 and S100A9 were generated by lentiviral transduction. Short-hairpin RNA (shRNA) vectors to S100A8 (TRCN0000053776) and S100A9 (TRCN0000053803) respectively cloned into the pRRLsin expression vector. were obtained from the Mission shRNA Library (Sigma-Aldrich). For viral production, these were cotransfected with packing vectors pMDLg/pRRE, rsv-REV and PVSV-g using the 3^rd^ generation lentiviral packing system.(30) The non-targeting hairpin SHC002 in pLKO.1-puro (CAACAAGATGAAGAGCACCAA) was included as a control. Selection was performed using 10 µg/ml puromycin (Sigma-Aldrich, Zwijndrecht, The Netherlands).

HT29 cells overexpressing tagged S100A8, S100A9 or both were generated by lentiviral transduction. For S100A8-HA, N-terminal 1 x HA-tagged S100A8 open reading frame was sub-cloned into an expression plasmid (pLV), under control of the CMV promotor, containing Blasticidin resistance. For S100A9-Flag, N-terminal 1x Flag tagged S100A9 open reading frame was sub-cloned into an expression plasmid (pLV), under control of the CMV promotor, containing Neomycin resistance. Lentivirus was produced in HEK293T cells using the third generation lentiviral packaging system as above.(30) Cells were selected using neomycin (750 µg/ml, G418, Invitrogen) or blasticidin (10 µg/ml Sigma-aldrich, R21001/11583677). For generation of S100A8-HA/S100A9-Flag expressing cells, S100A9-Flag expressing cells were subsequentially transduced using the S100A8-HA construct and selected using blasticidin (10 µg/ml).

### Cell culture experiments

Cells were plated in 24 well plates (1,0 x 10E5 per well).The next day cells were stimulated with 50 ng/ml rhIL17 (R&D,7955-IL/CF Lot DCVA1123021)), 50 ng/ml rhIL22 (Peprotech, 200-22, Lot#0710246 I1422), 10 ng/ml rhIFN-γ (R&D, 285-IF-100), 20 ng/ml rhTNF-α (Peprotech, #300-01-A Lot #031825 A2323) as indicated in figure legends.

### Digital spatial profiling

Geomix Digital Spatial Profiling was preformed according to manufacturer’s instructions and as previously described.(31) In brief, slides with 5 μM FPPE tissue sections were loaded on the Leica Bond RXm system for baking, deparaffinization and epitope retrieval in ER2 solution (Leica Biosystems, AR9640) for 20 minutes at 100°C. After incubation for 15 minutes at 37°C with 1μg/ml proteinase K (Thermo Fisher Scientific, AM2546) and post fixation in 10% NBF, the slides were incubated overnight at 37°C with the human Whole Transcriptome Atlas (WTA) probe panel. Followed by washing the slides in 50% formamide at 37°C, blocking and staining with antibody for pan-cytokeratin (AE1+AE3, Novus Biologicals) and nuclear SYTO13 staining (Nanostring, 121303303). After loading and scanning of the slides on the Geomx instrument, the various regions of interest were selected manually and a mask for epithelium (pan-cytokeratin positive) was generated to allow capture of the localized probes. Sequencing was performed on a NextSeq sequencer (Illumina).

### Analysis of DSP

DSP data was analysed using the established GeoMix workflows (GeomxTools, GeoMxWorkflows, both Nanostring) in R version 4.3.0 and R Studio 2023.03.1. Sequencing reads were converted to expression counts Preprocessing and QC was performed, using the following parameters: minSegmentReads = 1000, percentTrimmed = 50, percentStitched = 50, percentAligned = 30, percentSaturation = 50, minNegativeCount = 1, maxNTCCount = 1000, minNuclei = 50, gene detection rate > 5%. Analysis was limited to genes detected in a minimum of 5% of samples. Data was normalized using Q3 normalization, differential expression was calculated using t-test with False Discovery Rate (FDR) multiple testing correction. Anodermal samples were only included for reference purposes, and not included in the statistical analysis. Gene set enrichment analysis was performed using GSEA software (version 4.3.2) and genesets previously described (sets enhanced in fetal vs adult intestinal epithelium and enhanced in wound associated epithelium vs normal epithelium).(9, 32) Prediction of upstream activity was performed using Ingenuity Pathway Analysis software (IPA, Qiagen). Upstream regulators were restricted to endogenous compounds.

### Bulk RNA isolation and RNASeq

Tissue biopsies and curretage material were homogenized using the SilentCrusher M (Heidolph Instruments) in TRI-reagent (Sigma-Aldrich). Total RNA was extracted using the Bioline Isolate II RNA Mini Kit (Qiagen) according to manufacturer’s protocol. RNA quality was assessed using the Tapestation 4200 (Agilent) and RNA concentration was measured using the Qubit 2.0 Fluorometer (Invitrogen). cDNA libraries were generated using the KAPA RNA HyperPrep Kit with Oligo-dT enrichment (Roche) and sequencing was performed on the Illumina NovaSeq6000. Reads from files in the FASTQ format were mapped to the hg38 human reference genome using CLC genomic workbench (Qiagen) version 22.0.0.

Publicly available datasets were obtained from the GEO database (GSE117993 and GSE83245, only control subjects from both studies were included) and aligned using the same parameters. Comparative analyses were performed using the CLC genomic workbench after normalization and batch correction for either the sequencing run (curettage material) or the specific experiment (comparisons to public datasets). For analysis of differentially expressed genes, downweighing of outliers and filter on average expression for FDR correction was applied. Pathway analysis was performed by Gene Set Enrichment Analysis (GSEA and Fgsea packages in R).(33)

### Single cell RNASeq-Data generation

Samples were slowly thawed, removed from freezing medium and digested at 37 °C for 30 minutes using RMPI 1640 (52400-025, Thermo Fisher Scientific) supplemented with 0,5 mg/mL Collagenase D (11088866001, Roche), 4 ug/mL DNase I (11284932001, Roche) and 20ul/ml FCS. Digested tissues were filtered using a 100um cell strainer, washed, resuspended in phosphate buffered saline (PBS) containing 1% Bovine Serum Albumin (A3299-50ML; Sigma-Aldrich) and stained using 4’, 6-Diamidino-2-phenylindole-dihydrochloride (DAPI) (1:10.000, 18860.01, Serva) and anti-CD66b-Pe-Cy7 (1:20, 305104, eBioscience). Cell preparations were then sorted on a Flow cytometer (SH800S Cell Sorter) excluding CD66b+ (granulocytes) and DAPI-high (dead) cells. CD66b-DAPI-low cells were collected and concentrated prior to sequencing. 5,000 cells/sample were loaded to the 10X Chromium controller, encapsulated in lipid droplet, and cDNA and libraries generated using the 10x Genomics Single-Cell 3’ kit according to the manufacturers protocol (Chromium NextGEM Single Cell 3’). cDNA libraries were sequenced to using the Illumina Nextseq 500. Data was aligned against transciptome version GRCh38-3.0.0 using Cell Ranger software (10X Genomics) generating expression count matrices.

### Single cell RNASeq – Annotation

Unfiltered (raw) expression count matrices as obtained from Cell Ranger were further processed using the CLC Genomics Workbench. Samples were filtered for empty droplets and multiplets and normalized to obtain cellular expressions using the Single Cell Analysis Module. Cells were clustered by UMAP principle(34) and epithelial clusters where identified as those expressing EPCAM and/or CHD1. After filtering for excessive mitochondrial transcripts (cutoff <50%), cells expressing markers specific for other lineages were removed (PTPRC, CD19, MS4A1, CD3E, HBB, HBA1). Filtered expression matrix was exported and further analyzed using the Seurat package v4.(35–38) Epithelial cells were reclustered separately in order to increase resolution within the subpopulation, resulting in 7 subclusters. Specific expression was determined for each cluster compared to all other cells using non-parametric Wilcoxon rank sum test, with a minimum expression in 10% of cells Clusters were manually curated and assigned based on the most specific expression patterns (Enterocytes: *PHGR1, MUC12, PLA2G2A, LGALS4, PIGR*, Cycling: *MKI67, TOP2A, HMGB2, CENPF, NUSAP1*, Goblet: *SPINK4, MUC2, TFF3, FCGBP, ITLN1*, Enteroendocrine: *CHGA, SCG2, GRP, PCSK1N, CPE*, Tufts: *TRPM5, LRMP, FYB1, PSTPIP2, HPGDS*, WFDC2+: *KRT13, WFDC2, ANXA1, PSCA, S100A9*, Melanocyte-like: *DCT, MLANA, APOE, PMEL, IGFBP7*, Fig S3a).

### Single cell RNASeq – Pseudotime analysis

Pseudotime analysis was performed using the Monocle3 package.(39–41) Prior to pseudotime analysis, highly specialized/differentiated epithelial cell types (Enteroendocrine, Melanocyte-like, Tufts, Goblet) were excluded and remaining cells were reclustered to evaluate the developmental stages of enterocytes. Subclusters included early colonocytes (REG1A, RARRES2), colonocytes (KRT20, FABP1), late colonocytes (CEACAM1, BEST4), cycling epithelium (MKI67, TOP2A) and WFDC2+ (WFDC2, KRT13) Fig S4a). Differential expression between trajactories was determined by Regression analysis (cut off q_value<0.001). Pathway analysis was performed using the Fgsea package and the Gene Ontology pathway database.

### Immunohistochemistry

Paraffin embedded blocks were sectioned at 4.5um, deparaffinized with xylene, rehydrated and endogenous peroxidases were blocked with 0.01% H2O2 in PBS. Subsequently slides were boiled in 0.01 M sodium citrate buffer (pH 6) for 10 min at 120 °C for antigen retrieval and non-specific binding sites were blocked using PBS containing 1% bovine serum albumin and 0.1% Triton-X-100 (30 min, room temperature). Slides were incubated overnight with primary antibody diluted in the blocking buffer, anti-S100A9 (1:500; MAC387, Thermofisher). Brightvision anti-mouseHRP (ready-to-use; Immunologic, VWRKDPVM110HRP) was used as a secondary antibody. Antibody binding was visualized by adding chromagene substrate diaminobenzedine (DAB, Sigma-Aldrich), after which slides were counterstained using haematoxillin (Sigma-Aldrich) and dehydrated and mounted with Entellan (Sigma). Images were obtained using an Olympus BX51 microscope and quantified on a 0-4 intensity scale by a blinded investigator or using Fuji software. (42)

### RNA scope – in situ hybridization

In situ hybridization (ISH) was performed using RNAScope technology according to manufacturer’s protocol (Advanced Cell Diagnostics, ACD). In brief, slides were deparaffinized, treated with heat antigen retrieval and protease, followed by a series of hybridizations with the target-specific probe and amplifiers. The single-plex RNAscope probe against human CXCL17 (#513241) was used and detection was performed using the Fast RED reagent. Slides were counterstained with hematoxcylin, washed in PBS and mounted with ProLong Gold Antifade reagent with DAPI (Thermo Fisher Scientific).

### Reanalysis of CHIPSeq data

Public datasets were downloaded from the GEO database (GSE155421).(11) In the published study, MCF-10A–ER-Src cells were transfected using S100A8-Flag or S100A9-Flag and transformed using Tamoxifen. Crosslinking was performed, followed by nuclear sonification, immunoprecipitation using the Flag-tag and sequencing of the sheared DNA fragments.

Downloaded data was mapped against the GRCh38 genome. TF CHIP-Seq analysis was then performed using as foreground the triplicate tamoxifen treated S100A8-Flag or S100A9-Flag transfected samples with full IP and as background the input (ie non-precipitated) of the same samples. Cut-off for peak calling was set at p<0.05. Mapping to genes was performed against the GRCh38.107 gene mapping. All computational analysis was performed using the CLC Genomics Workbench (Qiagen).

### Chromatin Immunoprecipitation (CHIP) – qPCR

HT29 cells expressing both S100A9-FLAG and S100A8-HA were stimulated with the cytokine mix for 24 hours. After 24 hours cells were directly fixed by adding 250 x protein-protein-cross-linking solution (ChIP Cross-link Gold, Diagenode SA, Seraing, Belgium, cat.nr#C01019027) to culture medium for 30 min at RT and subsequent protein-DNA cross-linking with 1% Formaldehyde for 15 min at RT, which was quenched by adding of Glycin. Samples were further processed according to the iDEAL ChIP qPCR kit (Diaganode, cat.nr#01010180) protocol, with an optimized sonification protocol (7 cycles 30 s on/30 s off, in a Bioruptor Pico (Diagenode)), and subsequent immunoprecipitation using anti-FLAG or anti-HA beads. qPCR was performed on immunoprecipitated fractions and corresponding input samples and calculated as fraction of input sample, relative to control gDNA (PPIA) fractions, per gene (C3, CXCl17, LCN2, ACTB).

Primers for C3, CXCL17 and LCN2 were developed based on predicted peak regions from ref 15 (table S6).

### cDNA and quantitative PCR

cDNA was synthesized from 1ug of purified RNA using Revertaid reverse transcriptase according to protocol (Thermo Fisher Scientific, Landsmeer, The Netherlands) in the presence of both Oligo-dT primer and random hexamers primers. Quantitative real time PCR was performed using a SensiFAST SYBR No-ROX Kit (Waddinxveen, The Netherlands) according to the manufacturer’s protocol. Primer sequences are listed in Star methods. Optimal reference genes GAPDH and 36B4 were identified using the GeNorm algorithm and the geometric mean of these was used for normalization of data.(43) Data is shown as expression relative to the average of control.

### Immunoblotting and ELISA

Nuclear and cytoplasm extracts were isolated using the NE-PER kit (Thermo Fisher Scientific, #78833). For co-IP experiments freshly prepared nuclear extracts were immunoprecipitated using magnetic anti-HA-beads (Thermo Fisher Scientific, #88836) or magnetic anti-FLAG-M2-beads(Sigma-Aldrich, M8823) according to the manufacturer’s instructions.

Samples were run on SDS-PAGE gels under reducing conditions and transferred to nitrocellulose membranes (GE Health Care, Zeist, The Netherlands). Membranes were blocked by incubation in 5% non-fat milk (Nutricia, Wageningen, The Netherlands) in TBST (TBS + 0.1% Tween-20) for 2 hours at room temperature (RT) and subsequently incubated with S100A8 (1:200, Santa Cruz, Clone C-10, sc-48352, Lot *I1522), S100A9 (1:1000, CST, Clone D5O6O, 72590S, Lot #1), anti-NFĸB p65 (1:1000, CST, CloneD14E12, 8242P, Lot#4, anti-STAT3 (1:1000, CST, Clone 124H6, 9139S, Lot#10), anti-LaminB1-HRP (1:1000, CST, Clone D9V6H, 15068S, Lot#1) or β-Actin antibody (Sigma, Clone AC-15, A1978) antibodies in 2% milk/TBST overnight at 4ْ C. After incubation, membranes were washed 3 times with TBST, incubated with HRP conjugated secondary antibodies (Dakocytomation) in 2% milk/PBST for 2 hours at RT. Expression was detected by Lumilight Plus (Roche).

CXCL17 protein was detected in cell culture supernatants after 72 hour stimulation with the cytokine mix by ELISA, performed according to manufacturer (ThermoScientific, REF EH137 RB, Lot 2036060623), with overnight sample incubation at 4°C.

### Image analysis

Image analysis was performed by an observer blinded to the condition under evaluation. Scoring of S100A9 staining was performed on n=8 individual donors per group, using a 0-4 scale and nuclear and cytoplasmic staining was evaluated separately. Scoring of organoid morphology was performed on images obtained from 2 experiments using independent donors (each experiment n=9 images/condition). Individual organoids were designated as mainly round, containing significant budding or flattened by a blinded observer.

### Statistical analysis

Statistical analysis was performed using GraphPad Prism v9 or CLC Workbench and the relevant processing packages in R (for RNASeq data). Specific tests per experiment are denoted in the figure legends and included Mann-Whitney U test, Kruskal-Wallace with Dunn’s post-test, ANOVA with Dunnet’s post-test. Genome scale testing was corrected by False Discovery Rate (FDR). All tests were performed two sided. Data was considered significant with p<0.05.

### Public data accession

Public data used includes: Bulk RNASeq data, available from GSE117993 (pediatric samples) and GSE83245 (adult samples), CHIPSeq data: available from GSE155421 (CHIPSeq).

### Author contributions

MAB: Investigation, Formal analysis, Project administration, Writing of draft and revising, PJK: Conceptualization, Investigation, Formal analysis, Revision, SO: Resources, Investigation, Revision, SM: Investigation, Validation, MvR: Investigation, Project Administration, VM: Conceptualization, Investigation, Revision, JS: Investigation, Visualization, Revision, DAL: Investigation, Revision, WAB: Resources, Revision, GRD: Funding, Review, Conceptualization, CJB: Conceptualization, Resources, Project Administration, Funding, Supervision, Writing and Revision, MEW: Formal Analysis, Data curation, Funding, Supervision, Visualization, Writing and Revision.

## Acknowledgments

The authors wish to express their gratitude to S. Kenter and M. Jakobs (Core Facility Genomics, Amsterdam UMC) for help with the single cell experiments, I. Nijman and R. Geene (Useq facility, UMC Utrecht) for advice and assistance in the digital spatial profiling and T. Breit and M.J. Jonker (Molecular Analysis Department, Swammerdam Institute for Life Sciences, University of Amsterdam) for assistance in the bulk RNASeq experiments.

## Supplementary information

Supplementary figures-Figures S1-S6 and related legends

Supplementary Tables – Tables S1-S5, patient characteristics and S6, primer list

