## Supplementary figures and tables for "Nuclear Calprotectin mediates Epithelial Wound Healing Defects in Crohn’s Disease related Fistula"

**Becker at al. Supplementary information**

**Contents**

- **Supplementary figures S1-S6**
- **Supplementary tables S1-S6**

**Supplementary figures**

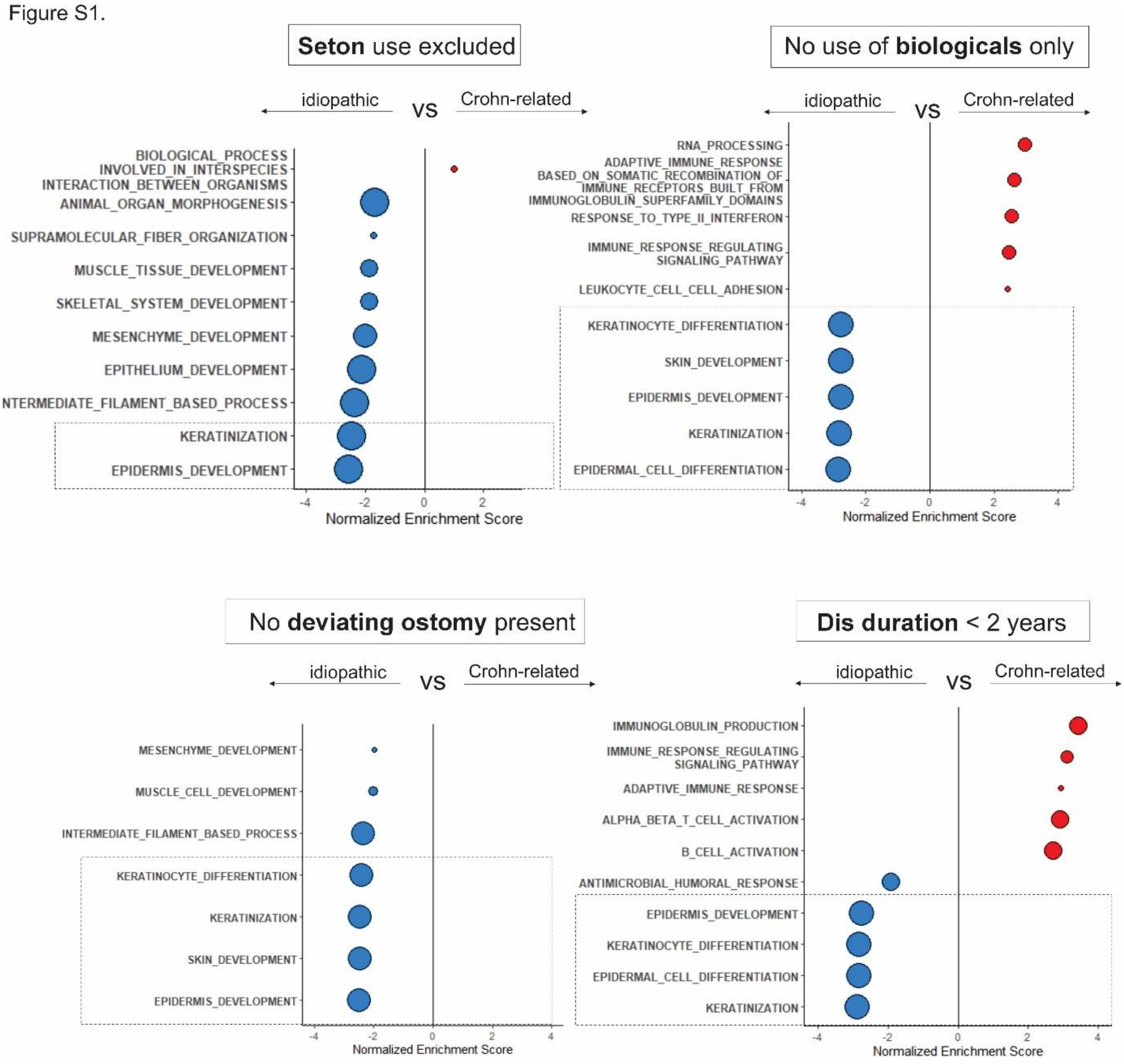


**Figure S1. No effect of IBD specific confounders on differentially activated pathways.** Differential gene expression and pathway analysis were performed excluding patients in whom a seton was present (A), only including those who did not receive any biological therapy (B), those without a deviating ostomy (C) or only those with a fistulizing disease duration of less than 2 years (D). In all cases, keratinization/(epi)dermal differentiation remained significantly enhanced in idiopathic fistula compared to Crohn-related fistula.

Figure S2

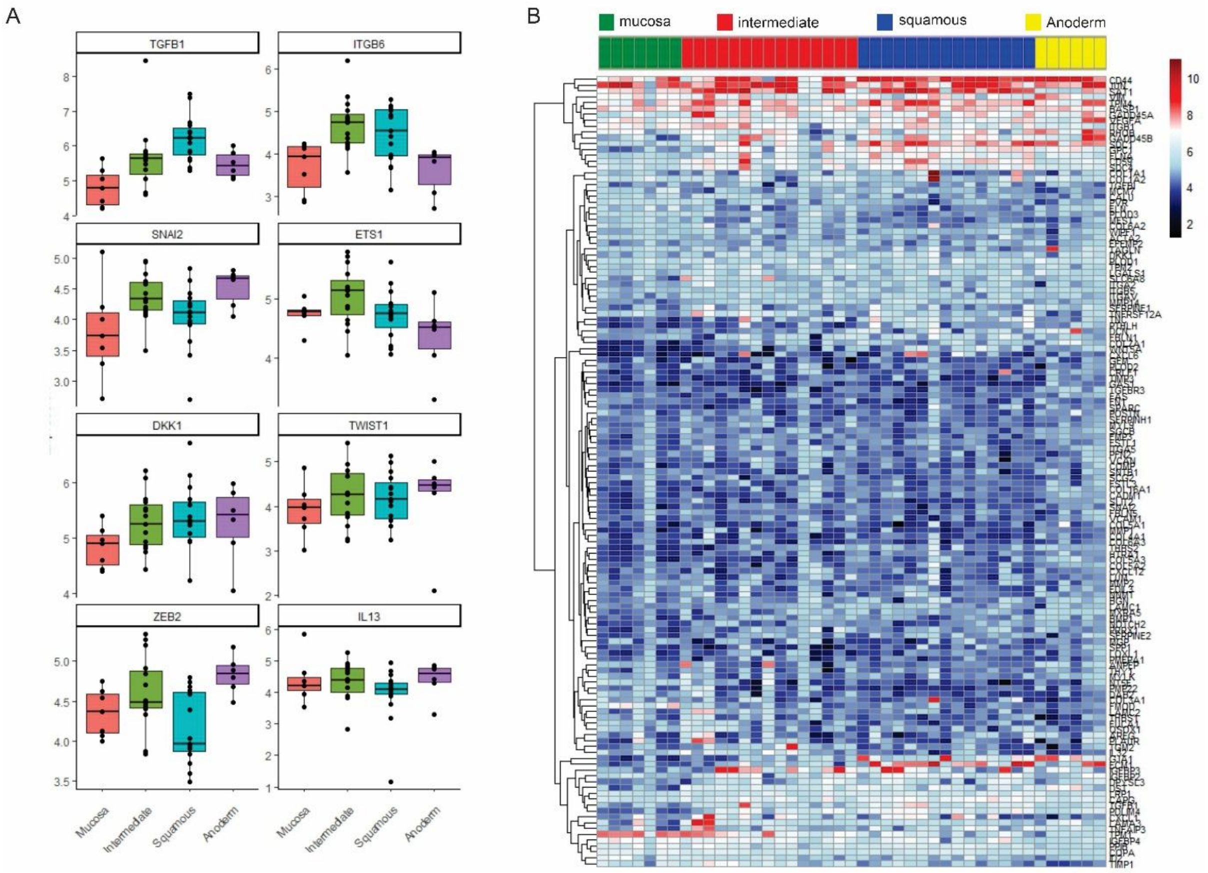


**Figure S2. Expression of EMT related transcripts over DSP samples.**

DSP analysis was performed on the mucosal (n=7), intermediate (n=15), squamous tissue (n=15) of the internal fistula openings, as well as anodermal samples (n=6) using a whole genome library. (A) Expression patterns over tissue subtypes of EMT markers previously associated with fistula formation. (B) Expression is shown for the genes included in the Hallmark geneset '*Epithelial to Mesenchymal Transition'* and passing quality control in the DSP analysis. Genes were clustered unsupervised.

Fig S3.

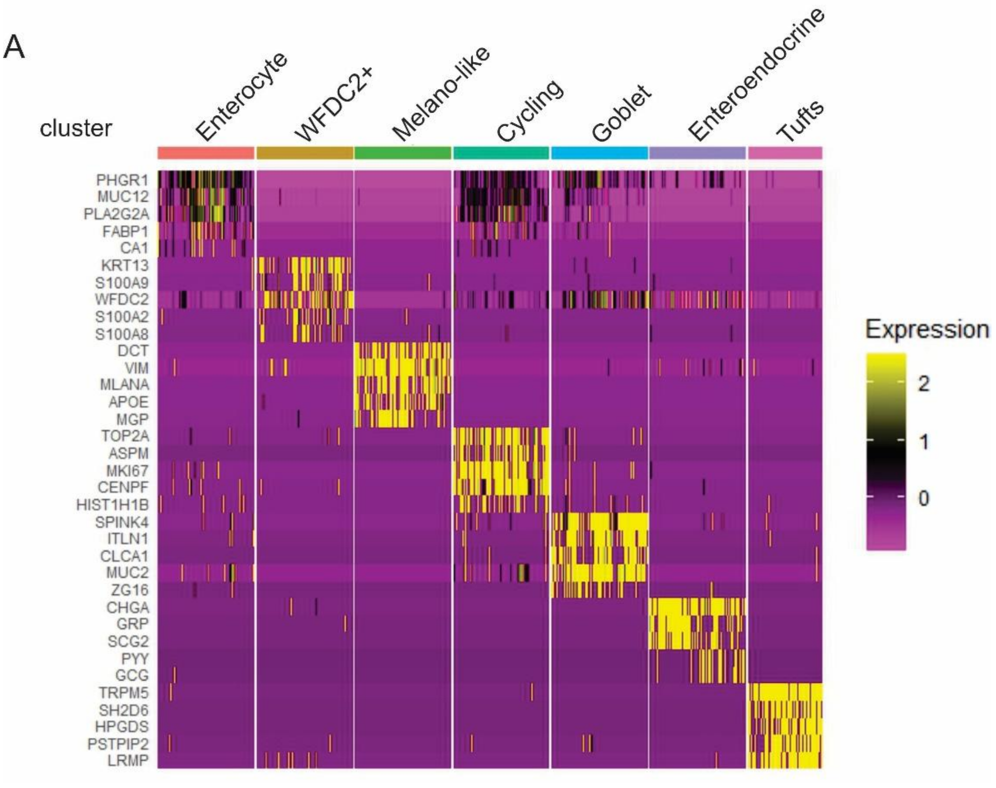


**Figure S3. Marker expression in single cell RNASeq clusters epithelium.**

(A) Expression of the top5 most specific makers per cluster of epithelial cells over all clusters. For visualization purposes, each cluster was downsampled to a maximum of 150 cells. (C) Expression profile of genes associated with the earlier described 'LDN' epithelial population across epithelial subsets.

Fig S4.

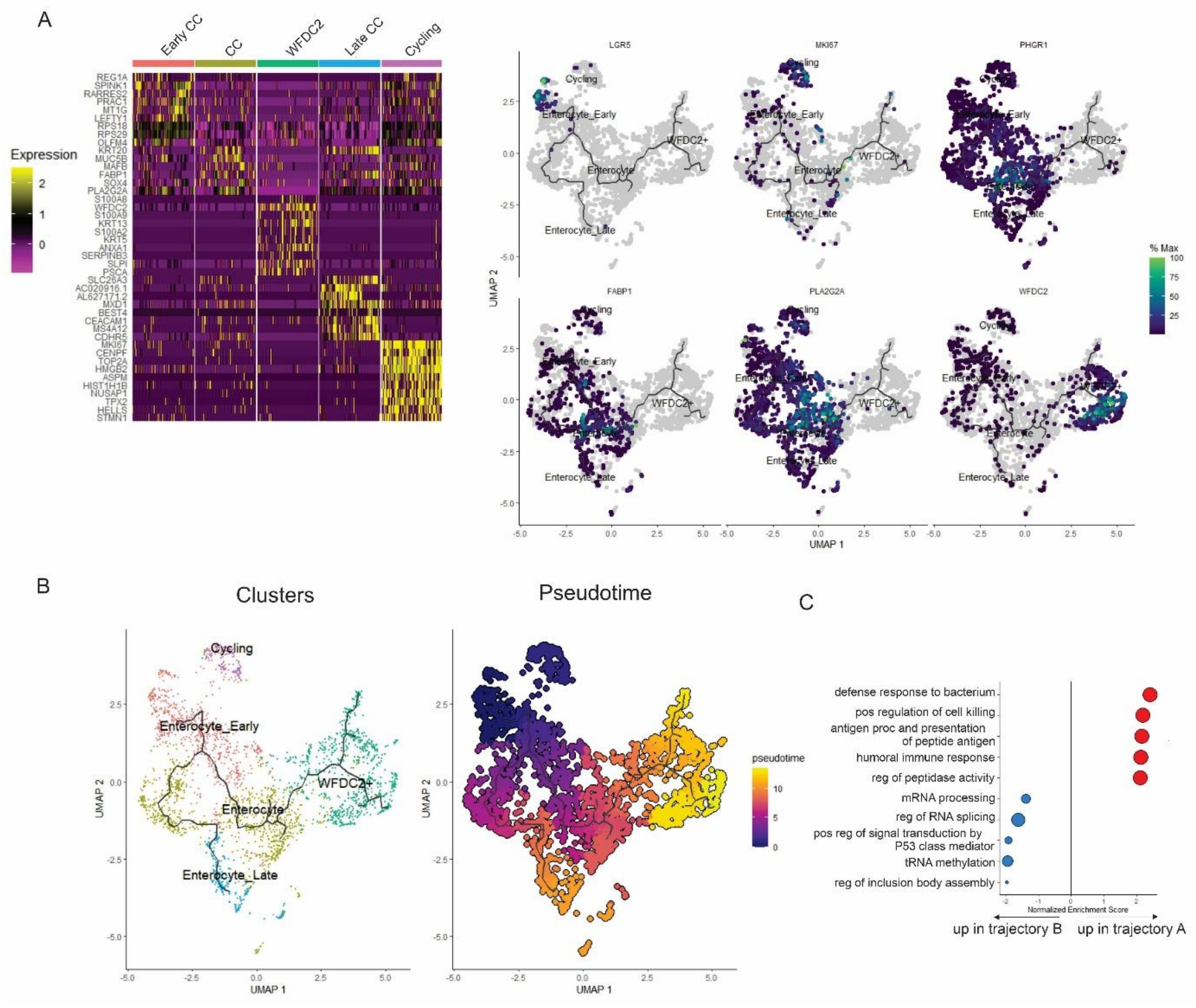


**Figure S4. Single cell analysis of internal fistula opening derived epithelium.**

(A) Expression of the top5 most specific makers per cluster of epithelial cells over clusters including differentiation of epithelium and after removal of highly specialized cells types. For visualization, each cluster was downsampled to a maximum of 150 cells. Right: clustered UMAP depicting selected markers. (B) Final annotated clusters and pseudotime analysis of the early enterocyte (RARRES2+, LGR5+) to late enterocyte and WFDC2+ cells.

Figure S5.

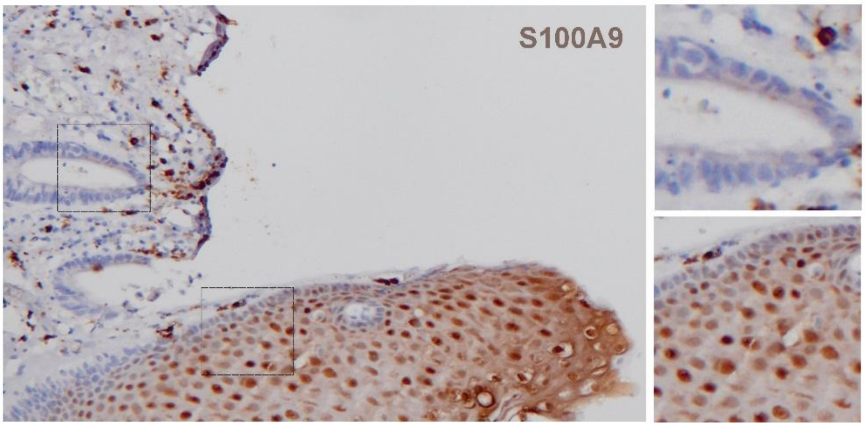


**Figure S5. S100A9 expression in rectal mucosa and squamous epithelium**.

Immunohistochemistry for S100A9 (brown) on the internal opening of a Crohn's related fistula indicates epithelial staining in the squamous regions but not in the adjacent intestinal mucosal area.

Figure S6.

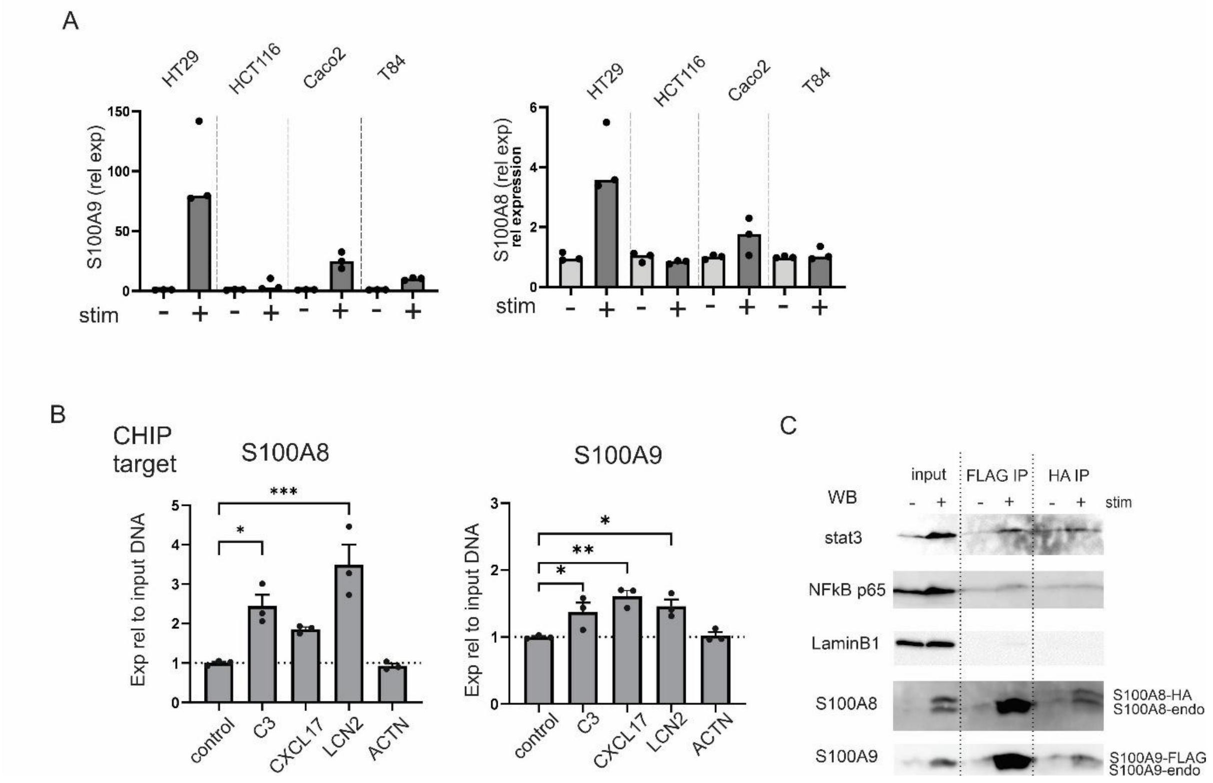


**Figure S6. Induction of S100A9 in intestinal and skin derived cell lines.**

(A) Intestinal cells (HT29) and keratinocyte derived cells (HaCat) were stimulated using a mixture of cytokines (IL17 50 ng/ml, IL22 50 ng/ml, IFN-γ 10 ng/ml, TNF-α 20 ng/ml)or IL17 alone for 24 hours. RNA expression is shown as relative to the unstimulated cells. Bars represent mean, Mann-Whitney U-test, * p<0.05. (B) Indicated cell lines were stimulated using a mixture of cytokines (IL17 50 ng/ml, IL22 50 ng/ml, IFN-γ 10 ng/ml, TNF-α 20 ng/ml) for 24 hours. RNA expression of is shown as relative to the unstimulated cells. Bar represent mean, error bars represent sem. (C) Chromatin immunoprecipitation was performed on S100A8-HA/S100A9-FLAG expressing HT29 cells stimulated with the cytokines above. Tagged S100A8 as S100A9 were used as IP targets, expression in resulting fractions was corrected for housekeeping gene PPIA and are shown as relative to the input DNA. Bars represent mean, individual dots represent repeat qpcr samples, One way ANOVA with Dunnets multiple testing correction, * p<0.05, ** p<0.01, *** p<0.001.

**Table S1.** Per patient information bulk RNASeq internal opening

| SampleID | Diagnosis | Gender | Age (yr) | Deviating s | Seton in sit | Proctitis du | Proctitis du | Biological u | Anti-TNF | Ustekinum | Vedolizum | Steroids sy | Thiopurines | Mesalazine |
|---|---|---|---|---|---|---|---|---|---|---|---|---|---|---|
| FI.126 | Crohn | male | 44 | No | Yes | No | No | No | No | No | No | No | Yes | No |
| FI.131 | Crohn | male | 67 | No | Yes | No | Yes | No | No | No | No | No | No | No |
| FI.132 | Crohn | male | 28 | No | Yes | No | No | No | No | No | No | No | No | No |
| FI.139 | Crohn | male | 75 | yes | Yes | Yes | Yes | No | No | No | No | No | No | No |
| FI.142 | Crohn | female | 38 | yes | No | No | Yes | yes | Yes | No | No | No | No | No |
| FI.148 | Crohn | male | 54 | No | No | No | Yes | yes | Yes | No | No | No | Yes | No |
| FI.149 | Crohn | female | 36 | No | No | No | Yes | No | No | No | No | No | No | No |
| FI.150 | Crohn | female | 22 | No | No | No | No | yes | Yes | No | No | No | Yes | No |
| FI.151 | Crohn | female | 36 | No | Yes | No | No | No | No | No | No | No | Yes | No |
| FI.152 | Crohn | male | 45 | No | Yes | No | No | yes | No | No | No | No | Yes | No |
| FI.157 | Crohn | female | 43 | No | No | No | No | yes | No | Yes | No | No | No | No |
| FI.158 | Crohn | female | 34 | No | No | Yes | Yes | yes | No | No | Yes | No | No | No |
| FI.177 | Crohn | male | 19 | No | No | No | No | No | No | No | No | No | Yes | No |
| FI.179 | Crohn | female | 48 | No | No | No | No | No | No | No | No | No | Yes | No |
| FI.180 | Crohn | female | 23 | No | Yes | No | Yes | yes | No | Yes | No | No | No | No |
| FI.182 | Crohn | female | 48 | No | Yes | No | No | yes | No | No | Yes | No | No | No |
| FI.185 | Crohn | male | 30 | No | Yes | Yes | Yes | yes | Yes | No | No | No | Yes | No |
| IRB.204 | Crohn | female | 38 | yes | No | Yes | Yes | Yes | No | No | Yes | No | No | No |
| IRB.212 | Crohn | male | 25 | No | Yes | No | No | No | No | No | No | No | Yes | No |
| IRB.288 | Crohn | male | 31 | No | No | No | No | No | No | No | No | No | Yes | No |

**Table S2.** Per patient information bulk RNASeq fistula tracts

| SampleID | Diagnosis | Gender | Age (yr) | eviating stoma | Seton in situ | octitis during surgery | Biological use | Anti-TNF | Ustekinumab | Vedolizumab | Steroids systemic | Thiopurines | Mesalazine |
|---|---|---|---|---|---|---|---|---|---|---|---|---|---|
| FI-100 | CD | Female | 39 | No | Yes | No | Yes | Yes | No | No | No | Yes | No |
| FI-102 | CD | Female | 57 | No | No | No | No | No | No | No | No | No | No |
| FI-103 | CD | Male | 29 | No | No | No | Yes | No | No | No | No | No | No |
| FI-107 | CD | Female | 28 | No | Yes | No | Yes | Yes | No | No | No | No | No |
| FI-111 | CD | Female | 26 | No | No | No | Yes | Yes | No | No | No | No | No |
| FI-113 | CD | Female | 40 | No | Yes | No | Yes | Yes | No | No | Yes | Yes | No |
| FI-115 | CD | Female | 19 | No | No | No | No | No | No | No | No | No | No |
| FI-116 | CD | Male | 75 | Yes | No | No | No | No | No | No | No | No | No |
| FI-117 | CD | Female | 33 | No | No | No | NA | NA | No | No | No | No | No |
| FI-120 | CD | Male | 21 | No | Yes | No | No | No | No | No | No | No | No |
| FI-124 | CD | Female | 35 | Yes | No | NA | Yes | No | Yes | No | No | No | No |
| FI-127 | CD | Male | 19 | No | Yes | NA | Yes | Yes | No | No | No | Yes | No |
| FI-128 | CD | Male | 33 | No | Yes | No | No | No | No | No | No | Yes | No |
| FI-131 | CD | Male | 67 | No | Yes | NA | No | No | No | No | No | No | No |
| FI-136 | CD | Male | 47 | No | No | No | Yes | Yes | No | No | No | No | No |
| FI-145 | CD | Female | 23 | Yes | Yes | Yes | Yes | Yes | No | No | No | No | No |
| FI-150 | CD | Female | 22 | No | No | No | Yes | Yes | No | No | No | Yes | No |
| FI-152 | CD | Male | 45 | No | Yes | No | No | No | No | No | No | Yes | No |
| FI-158 | CD | Female | 34 | No | No | No | Yes | No | No | Yes | No | No | No |
| FI-171 | CD | Male | 29 | No | No | No | Yes | Yes | No | No | No | Yes | No |
| FI-174 | CD | Female | 48 | No | Yes | No | Yes | No | No | Yes | No | No | No |
| FI-181 | CD | Female | 43 | No | Yes | No | Yes | No | No | No | No | No | No |
| FI-57 | CD | Male | 48 | No | Yes | No | Yes | No | Yes | No | No | Yes | Yes |
| FI-54 | CD | Male | 39 | No | No | Yes | Yes | No | Yes | No | No | No | No |
| FI-69 | CD | Male | 23 | No | No | No | Yes | Yes | No | No | No | Yes | No |
| FI-72 | CD | Female | 51 | No | No | No | Yes | No | Yes | No | No | No | No |
| FI-76 | CD | Female | 37 | No | Yes | No | No | No | No | No | No | No | No |
| FI-78 | CD | Male | 35 | No | No | No | No | No | No | No | No | No | No |
| FI-88 | CD | Female | 35 | No | No | No | Yes | Yes | No | No | No | No | No |
| FI-89 | CD | Female | 27 | No | Yes | No | Yes | Yes | No | No | No | No | No |
| FI-90 | CD | Female | 24 | Yes | Yes | No | Yes | Yes | No | No | No | Yes | No |
| IRB-212 | CD | Male | 25 | No | Yes | No | Yes | Yes | No | No | No | Yes | No |
| IRB-219 | CD | Female | 44 | No | Yes | No | No | No | No | No | No | No | Yes |
| IRB-222 | CD | Female | 40 | Yes | Yes | No | No | No | No | No | No | No | No |
| IRB-223 | CD | Male | 37 | No | Yes | No | Yes | Yes | No | No | No | Yes | No |
| IRB-227 | CD | Female | 60 | Yes | Yes | No | No | No | No | No | No | No | No |
| IRB-292 | CD | Female | 34 | Yes | No | Yes | Yes | Yes | No | No | No | Yes | No |
| IRB-298 | CD | Male | 32 | No | Yes | No | No | No | No | No | No | No | No |
| FI-109 | Crypto | Male | 36 | No | No | No | No | No | No | No | No | No | No |
| FI-129 | Crypto | Male | 49 | No | Yes | No | No | No | No | No | No | No | No |
| FI-176 | Crypto | Male | 51 | No | Yes | No | No | No | No | No | No | No | No |
| FI-178 | Crypto | Male | 62 | No | No | No | No | No | No | No | No | No | No |
| FI-43 | Crypto | Male | 51 | No | Yes | No | No | No | No | No | No | No | No |
| FI-44 | Crypto | Female | 62 | No | No | No | No | No | No | No | No | No | No |
| FI-45 | Crypto | Male | 68 | No | No | No | No | No | No | No | No | No | No |

| | | | | | | | | | | | | | |
|---|---|---|---|---|---|---|---|---|---|---|---|---|---|
| FI-48 | Crypto | Female | 45 | No | Yes | No | No | No | No | No | No | No | No |
| FI-60 | Crypto | Male | 41 | No | No | No | No | No | No | No | No | No | No |
| FI-64 | Crypto | Female | 43 | No | No | No | No | No | No | No | No | No | No |
| FI-65 | Crypto | Male | 62 | No | Yes | No | No | No | No | No | No | No | No |
| FI-85 | Crypto | Male | 50 | No | Yes | No | No | No | No | No | No | No | No |
| FI-95 | Crypto | Male | 55 | No | Yes | No | No | No | No | No | No | No | No |

**Table S3.** Per patient information digital spatial profiling

| SampleID | Diagnosis | Gender | Age (yr) | Deviating c | Seton in sit | Proctitis | Biologicals | anti-TNF | Ustekinum | Vedolizum | Tofacitinib | Steroids (s | Thiopurine | Mesalazine |
|---|---|---|---|---|---|---|---|---|---|---|---|---|---|---|
| FI120 | CD | Male | 21 | No | Yes | No | No | No | No | No | No | No | No | No |
| FI124 | CD | Female | 34 | Yes | No | No | Yes | No | Yes | No | No | No | No | No |
| FI57 | CD | Male | 47 | No | No | No | Yes | No | Yes | No | No | No | Yes | Yes |
| FI79 | CD | Female | 33 | No | Yes | No | No | No | No | No | No | No | No | No |
| FI87 | CD | Male | 40 | No | Yes | No | Yes | Yes | No | No | No | No | Yes | No |
| IRB190 | CD | Female | 61 | Yes | No | No | Yes | Yes | No | No | No | No | No | No |
| IRB204 | CD | Female | 38 | Yes | No | No | No | No | No | Yes | No | No | No | No |
| IRB269 | CD | Female | 36 | Yes | Yes | No | Yes | Yes | No | No | No | No | No | No |
| IRB273 | CG | Female | 60 | No | Yes | No | No | No | No | No | No | No | No | No |
| IRB295 | CD | Male | 33 | No | No | No | No | No | No | No | No | No | No | No |
| IRB274 | CD | Female | 27 | No | No | Yes | No | No | No | No | No | No | No | No |

**Table S4.** Per sample data for patients whose fistula tract was used in the organoids experiments

| SampleID | Gender | Age | Diagnosis | Medication at time of surgery | Type of surgery |
|---|---|---|---|---|---|
| FI395 | male | 53 | CD | none | colectomy |
| IRB 547 | male | 28 | CD | anti-TNF | colectomy |
| IRB533 | female | 27 | CD | anti-TNF, topical steroids | ICR |
| IRB510 | female | 56 | CD | anti-TNF | ICR |
| IRB524 | female | 53 | CD | anti-TNF, ustekinumab | ICR |
| IRB530 | male | 57 | CD | vedolizumab | colectomy |

**Table S5.** Per patient information patients included in the single cell RNASeq analysis

| Name | Disease | Gender | Age (yr) | Deviating d | Seton | Proctitis du | Proctitis ev | Biological u | Anti-TNF | Ustekinum | Vedolizum | Steroid sys | Steroid top | Thiopurine | Methotrexate | Mesalazine |
|---|---|---|---|---|---|---|---|---|---|---|---|---|---|---|---|---|
| FI.176 | CG | male | 53 | No | Yes | NA | NA | NA | No | No | No | No | No | No | No | No |
| FI.178 | CG | male | 64 | No | No | NA | NA | NA | No | No | No | No | No | No | No | No |
| FI.200 | CG | male | 70 | No | No | NA | NA | NA | No | No | No | Yes | No | No | No | No |
| FI.209 | CG | female | 36 | No | Yes | NA | NA | NA | No | No | No | No | No | No | No | No |
| FI.211 | CG | male | 40 | No | Yes | NA | NA | NA | No | No | No | No | No | No | No | No |
| FI.220 | CG | male | 56 | No | No | NA | NA | NA | No | No | No | No | No | No | No | No |
| FI.222 | CG | female | 37 | No | No | NA | NA | NA | No | No | No | No | No | No | No | No |
| FI.223 | CG | female | 32 | No | Yes | NA | NA | NA | No | No | No | No | No | No | No | No |
| FI.228 | CG | male | 58 | No | Yes | NA | NA | NA | No | No | No | No | No | No | No | No |
| FI.249 | CG | male | 43 | No | No | NA | NA | NA | No | No | No | No | No | No | No | No |
| FI.273 | CG | female | 53 | No | No | NA | NA | NA | No | No | No | No | No | No | No | No |
| FI.166 | CD | female | 20 | No | No | Yes | Yes | Yes | Yes | No | No | No | No | No | Yes | No |
| FI.168b | CD | female | 22 | No | Yes | No | No | Yes | Yes | No | No | No | No | No | No | No |
| FI.172 | CD | male | 33 | No | Yes | No | Yes | Yes | Yes | No | No | No | Yes | Yes | No | Yes |
| FI.175 | CD | female | 39 | No | Yes | No | No | Yes | No | Yes | No | No | No | No | No | No |
| FI.180 | CD | female | 25 | No | Yes | No | No | Yes | No | Yes | No | No | No | No | No | No |
| FI.183 | CD | female | 41 | No | Yes | Yes | Yes | Yes | Yes | No | No | No | No | No | Yes | No |
| FI.184 | CD | male | 47 | No | Yes | Yes | Yes | Yes | Yes | No | No | No | No | Yes | No | No |
| FI.185 | CD | male | 32 | No | Yes | Yes | Yes | Yes | Yes | No | No | No | No | Yes | No | No |
| FI.191 | CD | male | 33 | No | Yes | No | No | Yes | Yes | No | No | No | No | Yes | No | No |
| FI.198 | CD | female | 27 | No | No | No | No | No | Yes | No | No | No | No | Yes | No | No |
| FI.199 | CD | female | 38 | No | Yes | No | No | Yes | Yes | No | No | No | No | Yes | No | No |
| FI.205 | CD | female | 33 | No | Yes | No | No | Yes | Yes | No | No | No | No | Yes | No | No |
| FI.206 | CD | female | 41 | No | No | No | No | Yes | No | No | No | No | No | Yes | No | No |
| FI.210 | CD | female | 21 | No | Yes | No | Yes | Yes | No | Yes | No | No | No | No | No | No |
| FI.214 | CD | male | 37 | No | Yes | No | No | No | No | No | No | No | No | No | No | No |
| FI.236 | CD | male | 22 | No | Yes | No | No | Yes | Yes | No | No | No | No | No | No | No |
| FI.241 | CD | male | 35 | Yes | No | No | Yes | Yes | Yes | No | No | No | No | No | No | No |
| FI.245 | CD | female | 21 | No | No | No | No | Yes | Yes | No | No | No | No | No | No | No |
| FI.248 | CD | male | 26 | Yes | Yes | Yes | Yes | Yes | Yes | No | No | No | No | No | No | No |
| FI.268 | CD | male | 35 | No | No | Yes | Yes | Yes | Yes | No | Yes | No | No | No | No | No |

Table S6. Primer list for RT-qPCR

| **Target** | **FWD primer** | **REV primer** |
|---|---|---|
| C3 | GTGGAAATCCGAGCCGTTCTCT | GATGGTTACGGTCTGCTGGTGA |
| SLP1 | AGCGTGACTTGAAGTGTTGCATG | GAAAGGACCTGGACCACACAGA |
| PSCA | TGCTGTGCTACTCCTGCAAAGC | GAGTCATCCACGCAGTTCAAGC |
| TSPAN1 | TGCTGTGGTCGCCTTGGTGTAC | TGGTGAAGCCACAGCACTTGAG |
| CALML5 | GGAAACGGCACCATCAATGC | GTTTCCTTAGCTGGGCCTCC |
| LGALS2 | GACACAAGGTAGAAGGGGCAA | CTGAAGCGAGGGTTGAAATGC |
| MMP7 | AACGCTGGACGGATGGTAG | ATGAATGGATGTTCTGCCTGA |
| PPP1R1B | CCAACCCCTGTGCCTACAC | CCTCCATCTCTCTCGGACTC |
| WFDC2 | GCAAGAGTGCGTCTCGGA | TAATGTTCACCTGGGGGCAG |
| CXCL17 | ATGAAAGTTCTAATCTCTTCCCTC | CTACAAAGGCAGAGCAAAGCTTCTTAGC |
| S100A8 | TATCAGGAAAAAGGGTGCAGACG | TGCCACGCCCATCTTTATCA |
| S100A9 | GCAGCTGGAACGCAACATAG | TGTGTCCAGGTCCTCCATGA |
| LCN2 | GGTAGGCCTGGCAGGGAATG | CTTAATGTTGCCCAGCGTGAAC |
| KRT13 | AGGTGAAGATCCGTGACTGG | GTTGTTTTCAATGGTGGCG |
| GAPDH | AAGGTGAAGGTCGGAGTCAA | AATGAAGGGGTCATTGATGG |
| 36B4 | TCATCAACGGTACAAACGA | GCCTTGACCTTTTCAGCAAG |

Primer list for CHIP-qpcr

| Target | FWD primer | REV primer |
|---|---|---|
| PPIA | ATACGGGTCCTGGCATCTTG | AGGGCAACTTTTCATTTAGTCAG |
| C3 | CTGGAGAGGCGGTTTCTGA | CTGGAGAGCAAGCAGGTATTT |
| CXCL17 | CGCTGTTGTTGTTGCTGAA | TCTGTGCCCTTTCCAGTGTC |
| LCN2 | CTCCCCGTCCCTCTGTCTTG | CGCTGTGGTGGCTGCT |
| ACTB | TGGCAATGAGCGGTTC | GAAATGAGGGCAGGACTTAGC |
